# Multi-‘omics of host-microbiome interactions in short- and long-term Myalgic Encephalomyelitis/Chronic Fatigue Syndrome (ME/CFS)

**DOI:** 10.1101/2021.10.27.466150

**Authors:** Ruoyun Xiong, Courtney Gunter, Elizabeth Fleming, Suzanne D. Vernon, Lucinda Bateman, Derya Unutmaz, Julia Oh

**Affiliations:** The Jackson Laboratory, Farmington, Connecticut, USA; The University of Connecticut Health Center, Farmington, Connecticut, USA; Bateman Horne Center, Salt Lake City, Utah 84102

**Author notes:** Corresponding author and lead contact: Julia Oh, Ph.D., The Jackson Laboratory, 10 Discovery Drive, Farmington, CT.

## Abstract

Myalgic Encephalomyelitis/Chronic Fatigue Syndrome (ME/CFS) is a complex, multi-system, debilitating disability manifesting as severe fatigue and post-exertional malaise. The chronic dysfunctions in ME/CFS are increasingly recognized as significant health factors with potential parallels with ‘long COVID’. However, the etiology of ME/CFS remains elusive with limited high-resolution human studies. In addition, reliable biomarker-based diagnostics have not been well-established, but may assist in disease classification, particularly during different temporal phases of the disease. Here, we performed deep multi-‘omics (shotgun metagenomics of gut microbiota and plasma metabolomics) and clinical phenotyping of healthy controls (n=79) vs. two cohorts of ME/CFS patients – those with short-term disease (<4 years, n=75), and patients with long-term disease (>10y, n=79). Overall, ME/CFS was characterized by reduced gut microbiome diversity and richness with high heterogeneity, and depletion of sphingomyelins and short-chain fatty acids in the plasma. We found significant differences when stratifying by cohort; short-term ME/CFS was associated with more microbial dysbiosis, but long-term ME/CFS was associated with markedly more severe phenotypic and metabolic abnormalities. We identified a reduction in the gene-coding capacity (and relative abundance of butyrate producers) of microbial butyrate biosynthesis together with a reduction in the plasma concentration of butyrate, especially in the short-term group. Global co-association and detailed gene pathway correlation analyses linking the microbiome and metabolome identified additional potential biological mechanisms underlying host-microbiome interactions in ME/CFS, including bile acids and benzoate pathways. Finally, we built multiple state-of-the-art classifiers to identify microbes, microbial gene pathways, metabolites, and clinical features that individually or together, were most able to differentiate short or long-term MECFS, or MECFS vs. healthy controls. Taken together, our study presents the highest resolution, multi-cohort and multi-‘omics analysis to date, providing an important resource to facilitate mechanistic hypotheses of host-microbiome interactions in ME/CFS.

## Introduction

Myalgic Encephalomyelitis/Chronic Fatigue Syndrome (ME/CFS) is a complex, multi-system debilitating illness. The syndrome includes severe fatigue that is not alleviated by rest, post-exertional malaise (PEM), muscle and joint pain, headaches, sleep problems, hypersensitivity to sensory stimuli, and gastrointestinal symptoms (^1,2,3^). In the US alone, ME/CFS affects up to 2.5 million people (^4^). Our limited understanding of both the physiological changes associated with the syndrome and the underlying biological mechanisms are major impediments to identifying and developing both specific therapies and reliable biomarker-based diagnostics(^5,6^).

The human microbiome – the body’s trillions of bacteria, fungi, and viruses – has recently emerged as an important potential contributor to, or biomarker of ME/CFS (^7^). Patients have frequent gastrointestinal (GI) disturbances, and in lower-resolution studies based on 16S rRNA gene sequencing, altered gut microbiota (^7,8,9,10,11,12,13^). Compared to healthy controls, the microbial dysbiosis observed in ME/CFS patients was characterized by decreased bacterial diversity, overrepresentation of putative pro-inflammatory species, and reductions in putative anti-inflammatory species (^11,13^). However, sample sizes for these studies were relatively small with limited taxonomic resolution. Much remains underexplored vis a vis the potential functional consequences of ME/CFS-associated microbial changes.

An important function of the intestinal microbiota is metabolism (^14,15^), feeding both microbial and host processes in its dynamic, symbiotic, and mutualistic relationship with the host. For example, the metabolic products of gut microbiota can feed into host pathways as energy sources (^16^) or function as immune regulators (^17,18,19^), including short-chain fatty acids (^20^), metabolites of bile acids (^21^), or amino acid metabolites like tryptophan (^22^), respectively. Thus, the microbiome can modulate host physiology via direct stimulatory effects (^23,24^) or through secondary pathways coupled to metabolic processes (^25^). Such host-microbe interactions can be identified both through mechanistic studies (^26^) but also inferred by high resolution profiling (^27,28^) and integrated analyses of the gut microbiome (^29^) – the microbial fingerprint, and the host metabolome – the collective chemical fingerprint.

Here, we performed a high-resolution characterization of the gut microbiome and the plasma metabolome in two ME/CFS cohorts compared to healthy controls. In an important departure from current studies, we profiled a ‘short-term’ cohort (diagnosed within the previous four years), vs. a ‘long-term’ cohort (patients who have been suffering from ME/CFS for more than ten years). Our goal was to gain an understanding of the baseline molecular mechanisms by which changes in the ME/CFS microbiome may be reflected in circulating metabolic markers, which could then potentiate further alterations in host physiology. In addition, we sought to identify potential molecular and biological markers of ME/CFS progression between the short- and long-term cohorts. Finally, we collected detailed clinical and lifestyle survey metadata for association analysis. Shotgun sequencing of the fecal microbiome of 149 ME/CFS patients (74 short- and 75 long-term) vs. 79 age- and sex-matched healthy controls, we found that short-term ME/CFS patients had more significant microbial and gastrointestinal abnormalities, and that long-term patients tended to establish a stable, but individualized gut microbiome. However, long-term patients had significantly more irreversible health problems and progressive metabolic aberrations. Finally, integrating detailed clinical and lifestyle survey metadata with these high resolution ‘omics data allowed us to develop a highly accurate ME/CFS disease classifier. Taken together, our study provides a high resolution, multi-cohort and multi-‘omics analysis, providing new mechanistic hypotheses of host-microbiome interactions in ME/CFS.

## Results

### Data characteristics

We enrolled 228 participants; 149 with ME/CFS (74 ‘short-term’ and 75 ‘long-term’) and 79 approximately age- and sex-matched healthy controls (Figure 1, Table S1). The short-term and long-term cohorts were designed to obtain a better understanding of the biological processes during the progression of ME/CFS. The cohort was 96.5% Caucasian, with an average age of 43 years and 67% female, characteristics consistent with epidemiological reports that women are 3-4X more susceptible to ME/CFS than men (^30^). We collected detailed clinical metadata (Table S1, S2), stool samples for shotgun metagenomics, and blood for targeted metabolomic analysis. Clinical metadata (n = 228) and blood samples (n = 184) were collected at time of enrollment, followed by self-collection of fecal samples (n = 224) within the following two weeks (n=180 complete datasets of metadata, blood, and stool). We established a workflow to integrate the 906 clinical features into major disease markers to optimize data dimensionality (Figure 1). Whole-genome shotgun metagenomic sequencing of the stool samples generated an average of 10,801,733 high-quality and classifiable reads per sample, which were then reconstructed to examine gut microbiome composition (Table S3) and gene function (Table S5). Plasma was fractionated from blood and sent for targeted LC-MS analysis, where 1278 metabolites were identified for host molecular ‘omics profiling (Table S6). Finally, we analyzed each datatype individually, then altogether to build multi-‘omics models to describe and predict onset, stage of disorder, and associated microbial and metabolic features. This allowed us to target microbial pathways likely to affect host-microbiome interactions and alter the disease pathophysiology. For all datatypes, we performed two primary comparisons: 1) ME/CFS vs. healthy controls, to understand the broad differences inherent to the disease, and 2) short-vs. long-term ME/CFS vs. healthy controls, to understand disease progression.

**Figure 1.**
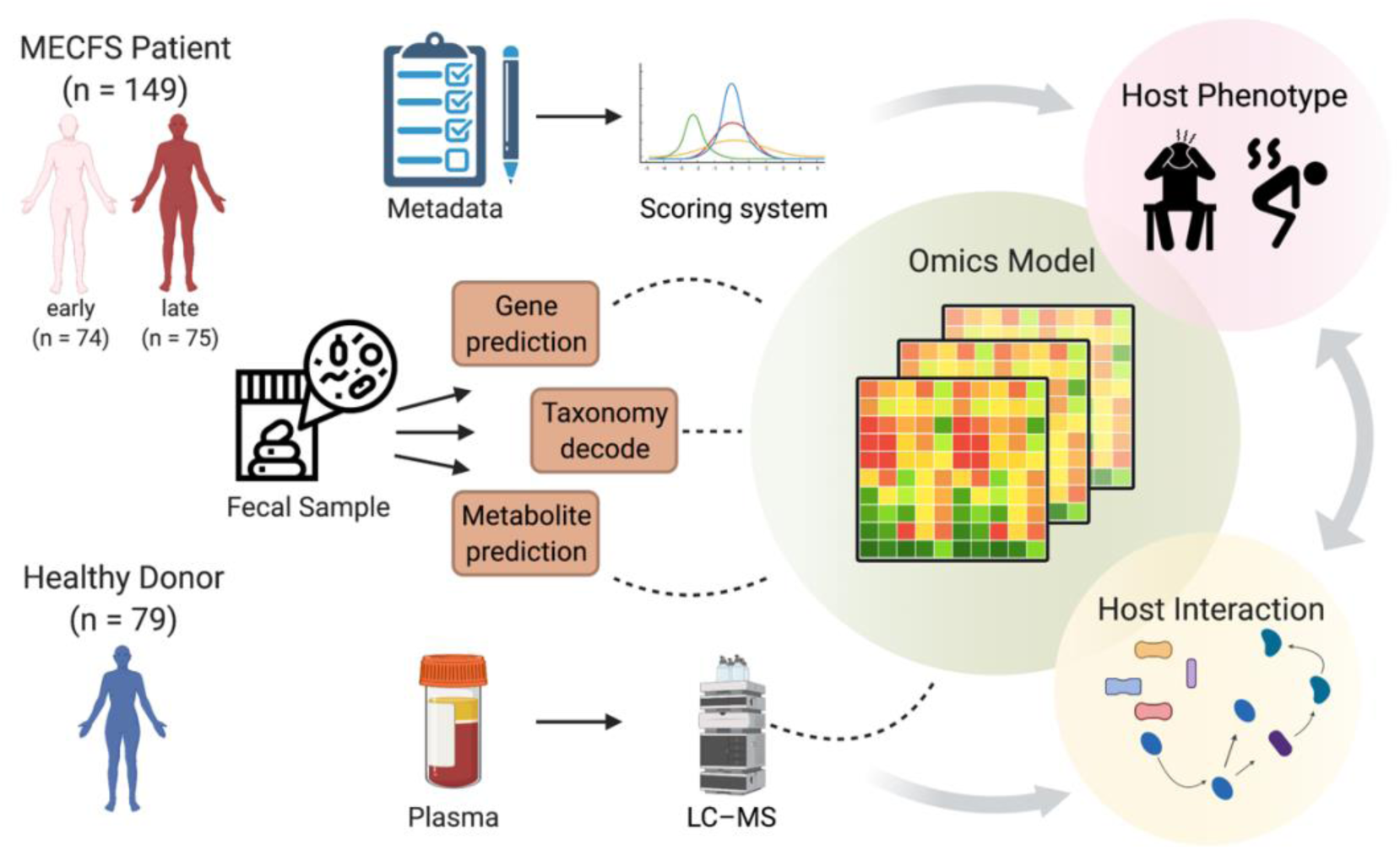
Summary of study design and analytical pipeline. We collected detailed clinical metadata, fecal samples, and blood samples for 228 individuals in three cohorts: healthy controls, patients with short-term (<4y) or long-term (>10y) ME/CFS. A comprehensive ‘omics workflow was constructed with the multi-data types (phenotypic, metagenomics, and metabolomics, respectively) and multi-computational models to understand potential host-microbe interactions. LC-MS, Liquid chromatography-mass spectrometry.

### The host phenotype in ME/CFS

To understand and interpret the host phenotype of ME/CFS compared to healthy controls, we collected comprehensive clinical metadata including detailed demographics and lifestyle information, an itemized dietary intake survey, medical history records, three general questionnaires regarding the physical and mental health of all participants, and five patient-specific surveys encompassing ME/CFS clinical symptoms and measurements (Table S2). In this analysis, we proposed a framework based on clinical know-how that ranked the contributions of various syndromes towards a composite score of disease severity. This framework allowed us to reduce the dimensionality of the clinical data for association analyses with ‘omics data and also to identify potential confounders.

We first excluded possible dietary biases, an important cofounder in microbiome (^31,32^) and metabolic studies (^33,34^). For example, we summarized dietary patterns, such as the frequency of meat/vegetable intake, etc., and then established that the dietary habits were comparable among all groups (Figure S1, Table S2). We also found that the frequency of previous acute infections, including mononucleosis (often caused by Epstein-Barr virus) and pneumonia were much higher in the patient cohorts (Figure S2, Chi-square test, p < 0.001), which supports the potential association of infections with the onset of ME/CFS (^35^). Our naïve Bayesian classification model (Figure S3A, area under the curve (a measure of classification accuracy), AUC = 0.85), which we used to identify clinical features that discriminate healthy controls vs. patients, showed that, besides some announced dysfunctions like orthostatic intolerance and fibromyalgia, most ME/CFS patients also suffered from additional complications, such as depression, headaches, constipation, and anxiety (^36,37^) (Figure S3B). Additionally, our scoring system, based on eight self-reported questionnaires (Table S2, see Method), showed that patients had significantly more anomalous mental and physical health conditions, as well as long-term poor life quality (Figure S3C, Wilcoxon rank-sum test, p < 0.001). Altogether, these high-resolution data echo the known pathophysiologies of ME/CFS and established the clinical characteristics of our cohorts (^38^).

### ME/CFS patients have decreased gut microbial diversity and greater heterogeneity

To begin to identify potential microbial mechanism(s) related to these symptoms, we first decoded the microbiota of ME/CFS patients. After classification of our shotgun metagenomic dataset to the species level (384 species passing quality cutoffs, see Method, Table S3), we examined community-wide metrics to understand if broad dysbiosis was observed in patients compared to controls. Clustering using principal coordinate analysis (Bray-Curtis dissimilarity distance, which reflects the similarity of microbiome composition between each pairwise set of samples) showed 1) most of sample variation was explained by the onset of ME/CFS (Figure 2F, permutational analysis of variance (PERMANOVA), p = 0.002) and was not influenced by the age difference between the control and patient groups (Figure S4), 2) patient samples had higher heterogeneity, as a population compared to controls (Figure 2G, Wilcoxon rank-sum test, p < 0.01). Interestingly, high heterogeneity was also observed in our recent study of frail older adults (^39^), suggesting a non-uniform adjustment to the host’s changing physiological conditions.

**Figure 2.**
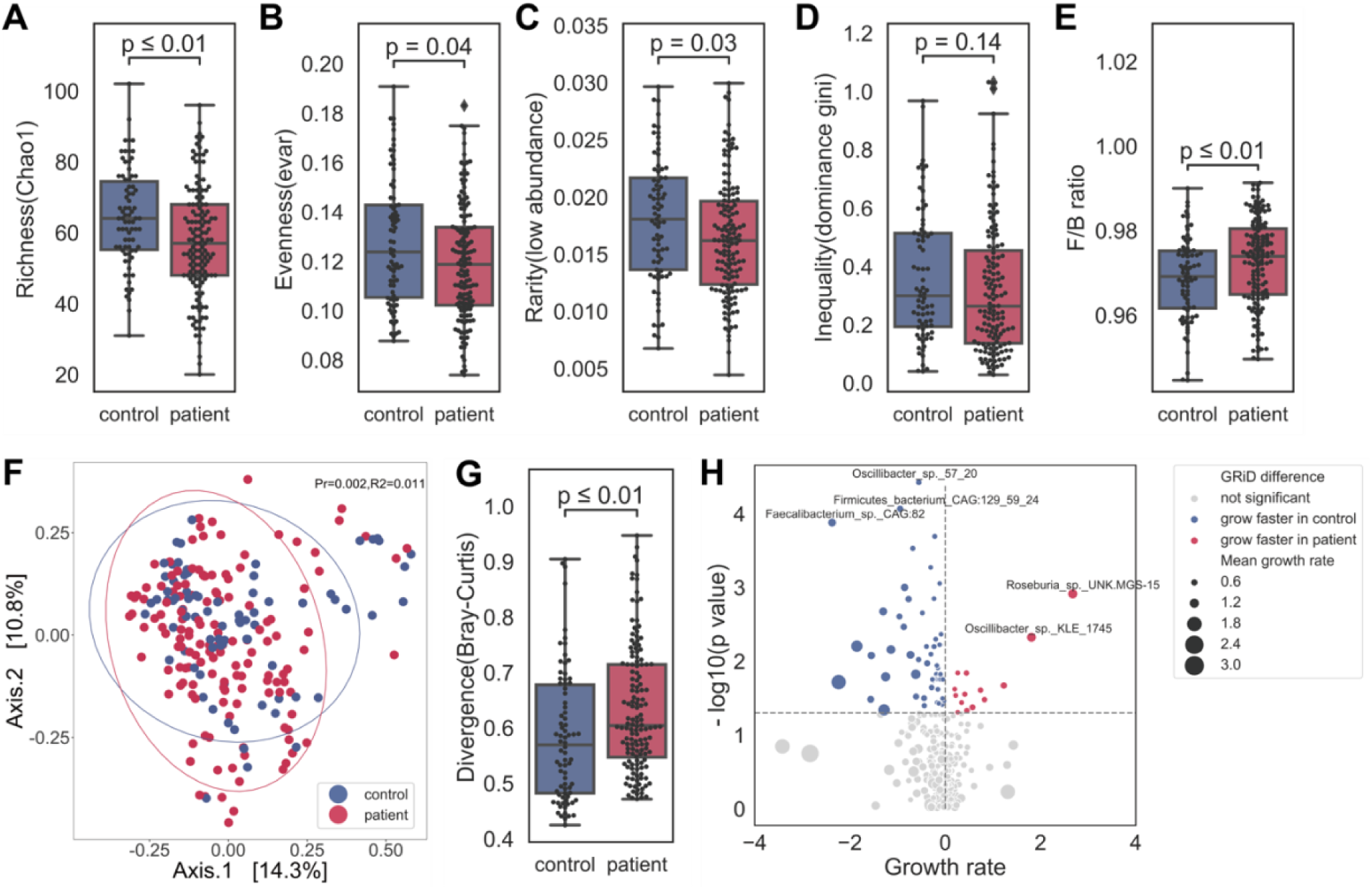
Microbial dysbiosis in ME/CFS is characterized by decreased diversity and greater heterogeneity. Comparing controls vs. ME/CFS patients (irrespective of disease stage), community structure differed in ME/CFS with A) decreased richness (Chao 1 index, which measures the number of observed species); B) decreased evenness (lower values of Smith and Wilson’s Evar index); C) increased rarity (larger proportion of the least abundant species (<0.2% relative abundance); D) increased inequality (larger Gini index of the dominant species (>0.2% relative abundance); E) decreased Firmicutes/Bacteroidetes ratio. p-values were computed by Wilcoxon rank-sum test. F) First and second principal coordinates of dimensionality reduction for Bray-Curtis dissimilarity distances, which measures pairwise similarity of two given samples). Values in brackets indicate the amount of total variability explained by each principal coordinates. p-value and R^2^ were calculated by permutational multivariate analysis of variance (PERMANOVA) test with patient/control as a variable. G) Increased heterogeneity observed in ME/CFS as measured by divergence, or Bray-Curtis dissimilarity. p-value was computed by Wilcoxon rank-sum test. H) Volcano plot showing differences in predicted growth rate in select species in ME/CFS. Each dot indicates a microbe, sized by the value of its inferred growth rate (Table S4). The x-axis shows the absolute difference (mean growth rate in patient – mean growth rate in control) and the y-axis is the log10(p-value, Wilcoxon rank-sum test). Species that were predicted to grow faster in patients were colored red and slower in blue. p-value > 0.05 was considered not significant (gray).

High microbial biodiversity has increasingly been associated with ecosystem health (^40^), with a less diverse (fewer members) and less even structure (i.e., more heavily weighted with fewer members) associated with decreased resilience and susceptibility to pathogenic colonization (^41^). ME/CFS patients had fewer community members (Figure 2A, Chao 1 index; Wilcoxon rank-sum test, p <0.001), and lower Evar evenness (Figure 2B, Wilcoxon rank-sum test, p <0.05). To understand if there were specific species lacking in patients, we calculated a rarity and dominance index. A decrease of Chao 1(Figure 2A) and rarity (Figure 2C, Wilcoxon rank-sum test, p <0.05) indicated that ME/CFS patients lost more low abundance members. The higher Gini index from those dominant species implied greater inequality (Figure 2D, Wilcoxon rank-sum test, p <0.01), with highly abundant commensal species, such as *Bacteroidaceae* members *Bacteroides (B*.*) vulgatus, B. uniformis, B. stercoris*, occupying a much larger portion of the population.

Next, we noted a modest change in the ratio of the overall relative abundance of Firmicutes to Bacteroidetes phyla (Figure 2E). This ratio has been implicated in some chronic disorders and inflammatory diseases such as obesity, diabetes, and inflammatory bowel disease (IBD) (^42,43^) and is of interest as a potential contributor to the immune dysbiosis observed in ME/CFS(^44^). At the family level, ME/CFS patients had reduced relative abundance of *Oscillospiraceae* (p < 0.001) and *Odoribacteracea* (p = 0.015), both reported in other chronic inflammatory disorders (^45^), and had significantly elevated *Clostridiaceae* (p < 0.001) and *Bacteroidaceae* (p = 0.033, Wilcoxon rank-sum test), enriched in IBD(^46^) and type I diabetes(^47^), respectively.

Finally, we note that metagenomic analyses reconstruct gene-coding potential but do not reflect the dynamic nature of the microbiome. We thus predicted in-situ growth rate of individual microbes with our Growth Rate InDex (GRiD) algorithm, which leverages coverage differences over a microbe’s genome to infer microbial growth rate, a proxy for metabolic activity in a community (^48^). Seventy-two species had significantly different growth rates in ME/CFS patients compared to controls. 14 replicated faster, including *Oscillibacter_sp*.*_KLE_1745* and *Roseburia_sp*.*_UNK*.*MGS-15*, and 58 more slowly, including *Oscillibacter_sp*.*_57_20, Firmicutes_bacterium_CAG:129_59_24*, and *Faecalibacterium_sp*.*_CAG:82* (Figure 2H, Table S4). Interestingly, the species that grew slower in ME/CFS were mostly Firmicutes, suggesting a further nuance to the already disproportionate Firmicutes:Bacteroides ratio. Taken together, the gut microbiome of ME/CFS, like in aging and other chronic inflammatory disorders, was characterized by modest but broad dysbiosis, including a less diverse and more uneven gut microbiome community with higher heterogeneity and altered Firmicutes:Bacteroidetes ratio, supported by a group of slower-replicating Firmicutes species.

### Microbial, metabolic, and genetic biomarkers of ME/CFS

Currently, there are no approved laboratory diagnostics available for ME/CFS (^6^), likely due to the heterogeneity of the disease. We hypothesized that the combination of environmental and clinical factors may assist in a more comprehensive disease classification. To determine the potential of metagenomic and metabolomic features to improve classification of ME/CFS, we constructed multiple state-of-the-art classifiers for deep profiling. For metagenomic data, we used both microbial species as well as gene relative abundance (identified by KEGG gene profiling, see Methods, Table S5) for our classifier, as we hypothesized that both species relative abundance as well as gene-level differences could differ between cohorts. Thus, we constructed four models based on 1) species and 2) KEGG gene relative abundances, 3) normalized abundance of plasma metabolites or 4) a combination of all three (multi-‘omics, see Methods), in addition to the clinical classifier described above. Irrespective of the model, results obtained using multi-‘omics data (gradient boosting model, AUC=0.90) outperformed any individual dataset, followed by the metabolome (AUC = 0.82), KEGG gene profile (AUC = 0.73), and species relative abundance (AUC = 0.73, gradient boosting model, Figure 3B; LASSO logistic, SVM, and Random Forest, Figure S5). This improved performance of multi-‘omics for differentiating ME/CFS patients from healthy controls underscores the complementarity of different ‘omics in describing the molecular processes that can occur with shifts in the host physiological state.

**Figure 3.**
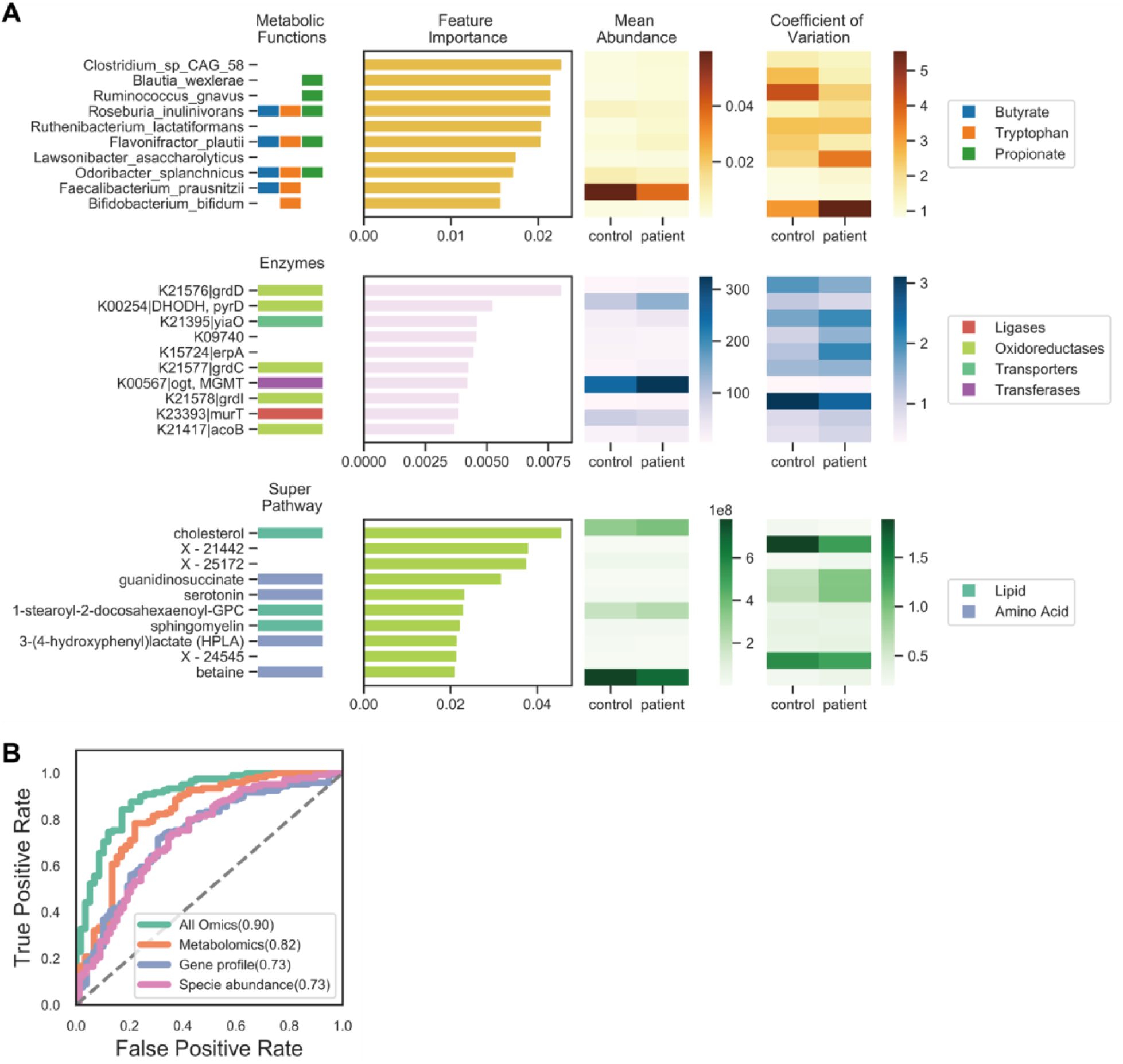
Out-performing multi-‘omics model identifies microbial, metagenomic, and metabolic biomarkers for ME/CFS compared to controls. A) Biomarkers from three supervised Gradient Boosting (GDBT) models are shown. Models from top to bottom: species relative abundance, relative abundance of KEGG gene profile, normalized abundance of plasma metabolomics. The top ten most important features in each model are shown together with their general functional class, raw abundance, and variance. From left to right: 1. Functional annotations: species relative abundance model - the metabolic function (capacity of butyrate, tryptophan, and propionate pathway); KEGG gene profile model - the class identification of the enzyme; metabolomics models - the superfamily for the metabolite; 2. Feature importance: features were ranked by their contribution to the model on the y-axis; the x-axis indicates the feature importance value from each model; 3. Average feature abundance in control and patient groups (Figure S5); 4. Variation in mean relative abundance in control and patient groups with coefficient of variation. B) Performance of the classifiers using area under the curve (AUC) was evaluated using 10 randomized and 10-fold cross-validations for each model: species relative abundance (pink), KEGG gene profile (blue) or metabolites (orange) alone, or taken altogether (‘omics, green), which used the combination of the top 30 features from three models.

We then examined the most discriminatory features for each of the individual ‘omics models to identify potential biomarkers and to further biological interpretation (Figure 3A, S5, Table S7). Low abundance microbes, i.e., those with relative abundance <1%, comprised the top 10 most discriminatory features. Among these were five microbes implicated in tryptophan metabolism, four of which were significantly reduced in ME/CFS patients (*Faecalibacterium (F*.*) prausnitzii, Odoribacter splanchnicus, Roseburia inulinivorans, B. bifidum*). Four butyrate producers were predicted, of which *F. prausnitzii*, an abundant gut commensal(^49,50,51^), was significantly decreased in ME/CFS (Wilcoxon rank-sum test, p < 0.01, Figure S5). Metabolites of both pathways (butyrate examined extensively later) are key immunomodulatory molecules that play pivotal roles in the regulation of metabolic and endocrine functions(^22,52–55^). Thus, an overall reduction in these microbes might potentiate the ME/CFS disease process. In addition, microbes capable of microbial fermentation to produce propionic acid, which can negatively influence growth of other microbes, were discriminatory for ME/CFS patients (increased relative abundance of *B. wexlerae, fermentation gnavus, Flavonifractor plautii*, and decreased *F. prausnitzii, B. bifidum*). Examining discriminatory KEGG gene features, we found 3 genes of interest had decreased relative abundance in ME/CFS patients. *grdD, grdE*, and *grdI* constitute the betaine reductase complex component, which produces betaine, an important anti-inflammatory metabolite(^56,57^). This aligned with discriminatory metabolomic markers, in which betaine was also identified as discriminatory, with a reduced concentration in patient plasma compared to controls. Other metabolic biomarkers were primarily in the lipid or amino acid super families. Besides betaine, sphingomyelin, serotonin, and cholesterol were highly discriminatory features, and have also been previously reported to be altered in ME/CFS patients (^6,58,59,60^).

### Microbial dysbiosis occurs in short-term ME/CFS and stabilizes in long-term disease

Our analyses thus far investigated major differentiating features between ME/CFS patients and healthy controls. We then wondered if microbial and metabolomic markers changed significantly during the progression of ME/CFS, or if early features could predict later severity. We then analyzed our dataset comparing the short-term to the long-term group. Because the age of the long-term patients was higher (Table S1, mean ± standard deviation (SD) 47± 1.4 years vs. 42± 1.5 and 40± 1.6 for the short-term group or healthy controls, respectively), we first evaluated the effect of age as a confounder but found no significant differences (PERMANOVA with age as a variate, p > 0.05, Wilcoxon rank-sum test, p > 0.05 (≤50 years old (yo) vs. >50 yo subgroups, Figure S4).

We then examined overall microbial composition differences between the patient cohorts as previously performed (Figure 4A). Interestingly, differences in the gut microbiome were more pronounced and variable during early stages of the disease based on the Chao1 index(p < 0.05), Evar evenness(p < 0.01), and Bray divergence (heterogeneity, p < 0.05) compared to long-term and controls (Figure 4C, 4D, 4F, Wilcoxon-rank sum test). As noted, heterogeneity was a feature we observed associated with aging, particularly frail aging(^39^). As we found no effect of age here, we conjecture that these differences were driven by ME/CFS disease duration.

**Figure 4.**
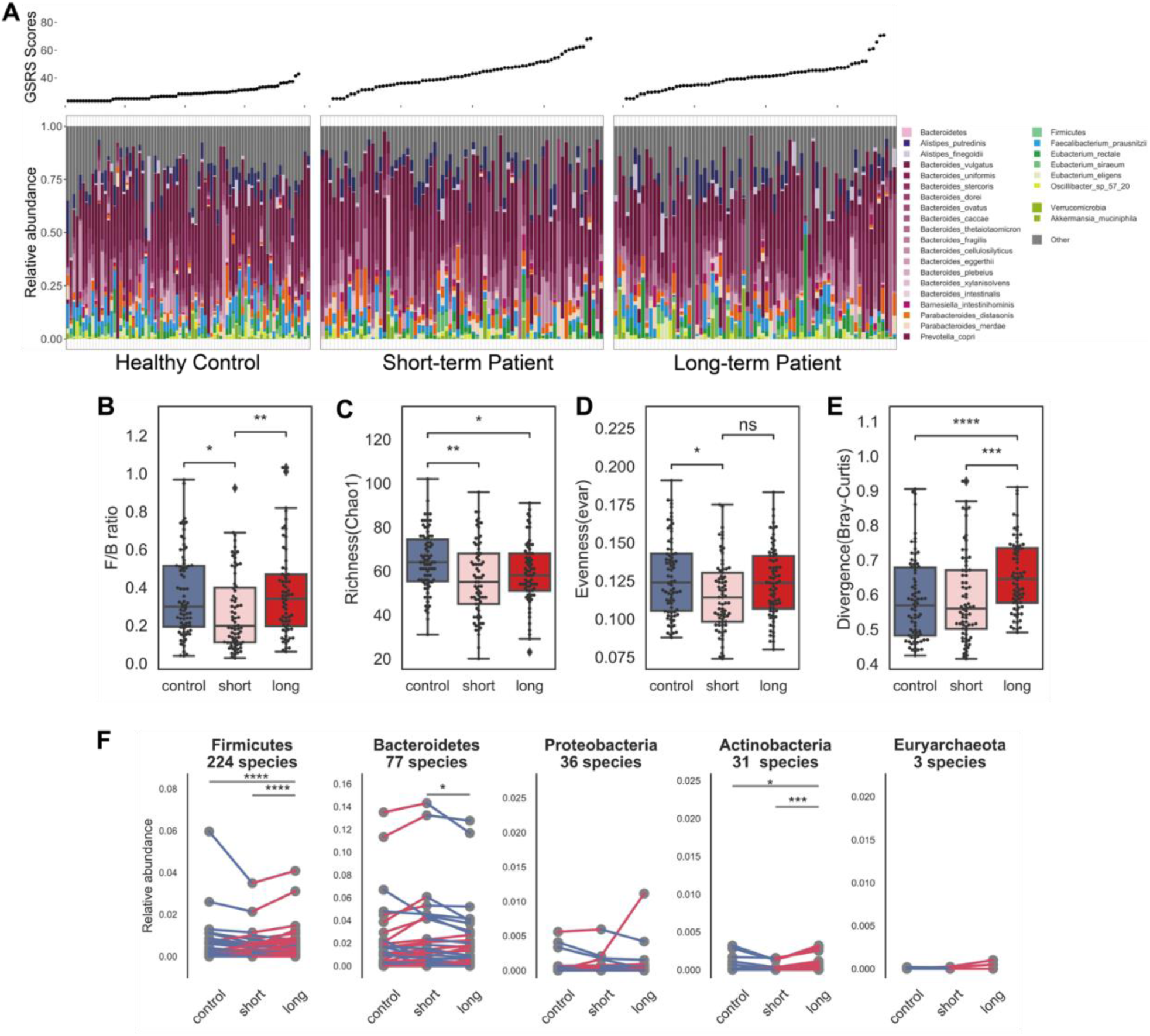
Significant microbial dysbiosis in observed in short-term ME/CFS. A) Taxonomic classification at species-level resolution for all individuals in the three cohorts: healthy controls, short-term patients, and long-term patients. Relative abundances of the most abundant gut species (top 25) are presented with gray representing the aggregate relative abundance of the remaining species. Gastrointestinal Symptom Rating Scale (GSRS) score, indicating the scale of gastrointestinal abnormality, is shown above for each individual. Similarly to Figure 2, we showed differences in the microbial community structures in short-term and long-term patients in: B) composition at phylum-level – decreased Firmicutes/Bacteroidetes ratio in short-term patients; C) reduced richness in short- and long-term (Chao 1 index, which measures the number of observed species); D) decreased evenness in short-term (lower values of Smith and Wilson’s Evar index). p-values were computed by Wilcoxon rank-sum test. E) Increased heterogeneity observed in long-term cohort as measured by divergence, or Bray-Curtis dissimilarity. p-value was computed by Wilcoxon rank-sum test. F) Each species in the five most abundant phyla were compared among three groups to observe the dynamics of gut community with respect to progression of disease. Here, each point represents the average relative abundance for a given species, connected with a line that is colored by increase (red) or decrease (blue). p-values were computed by Wilcoxon signed-rank test. p-value annotation legend: ns: p > 0.05, *: 0.01 < p <= 0.05, **: 0.001 < p <= 0.01, ***: 1e-04 < p <= 0.001, ****: p <= 1e-04.

At the taxonomic level, the differences observed in ME/CFS patients vs. controls were similarly more pronounced in short-term patients. A significant shift in the *Bacteroidetes* (77.5%) to *Firmicutes* (19.4%) ratio was observed only in short-term but not long-term patients (Figure 4B, Wilcoxon-rank sum test, p < 0.05 short-term; p > 0.1 long-term vs. controls), explaining the modest difference previously observed (Figure 2E). Comparison of the species relative abundance between the three cohorts showed that in the short-term group, the loss of diversity was driven by a relative reduction in low abundance bacteria that were more predominant in the long-term patients, with common commensal members like *Bacteroides* (Wilcoxon-rank sum test, p < 0.05) and *Parabacteroides* (Wilcoxon rank-sum test, p < 0.01) dominating. A decrease in *Prevotella* and *Faecalibacterium*, both of which have characterized anti-inflammatory functions (^61,62^), were observed in both short-term and long-term patients compared to healthy controls.

To further resolve species-level differences between short-term and long-term patients, we performed a pairwise comparison for the mean relative abundance of every species in the top five most abundant phyla (Figure 4F). We discovered that species that had atypical relative abundance in short-term patients with respect to healthy controls tended to stabilize in late-stage patients, i.e., become relatively closer to the relative abundance observed in controls. For example, most Bacteroides species, especially highly abundant ones, were greatly increased in the short-term group (Wilcoxon signed-rank test, p < 0.001) then slightly decreased in the long-term group (p < 0.001). Meanwhile, Firmicutes and Actinobacteria species decreased in both the short-term (Wilcoxon signed-rank test, p < 0.001) and the long-term groups (p < 0.001). However, in the long-term group, the magnitudes of relative abundances of Bacteroides, Firmicutes and Actinobacteria were relatively closer to those found in healthy individuals.

Taken together, we found that microbial dysbiosis is most marked early in ME/CFS disease, characterized by a loss of diversity driven by low abundance bacteria with high heterogeneity. Over time, we conjecture that the microbiota then stabilizes and reverts to an ecosystem more characteristic of healthy controls, with the reacquisition of some low abundance species and normalization of diversity. These results highlight the importance of stratifying by disease duration to identify features relevant to disease progression.

### Severe phenotypic and metabolic abnormalities in long-term ME/CFS

As described previously, we next constructed individual and multi-‘omics models to differentiate disease durations and controls. We followed this by over representation analysis (ORA) on metabolic pathways and Bayesian classification on phenotypic abnormalities to further pinpoint distinctive patterns in different stages of ME/CFS (see Methods). Overall classification accuracy was lower than for overall disease but still relatively accurate leveraging multi-‘omics data (AUC=0.82, gradient boosting model, Figure S6). Low (<0.5%) relative abundance bacteria, including several putative butyrate producers (*Clostridium* sp.) (^63^), were identified as potential biomarkers. The metabolomics model also identified two cholesterol and several lipid metabolites as discriminatory (Figure 5A, Table S7). This was consistent with the phenotypic classifier which identified many metabolic-related abnormalities, including high LDL cholesterol or triglycerides(^64,65^), hypothyroidism(^66^), obesity(^67^), hypoglycemia(^68^), and anecdotally reported complications, like constipation(^69^), sleep disturbances(^70^), and endometriosis(^71^) to be more predictive in long-term ME/CFS. However, we note that some of these phenotypes also can increase with age, which is largely matched in our cohort but may contribute to these differences (^72,73,74^). In addition, we found that fibromyalgia(^75^) was a key feature distinguishing short-vs. long-term disease, as was a trend towards worsening sleep problems and more pronounced post-exertional malaise in the later stage of the disease (Figure 5B). Interestingly, we identified poor appetite(^76^) to be the most distinguishable phenotype in the short-term group, which is consistent with a trend of more GI disturbances in the early stage (Figure 5B, 4A).

**Figure 5.**
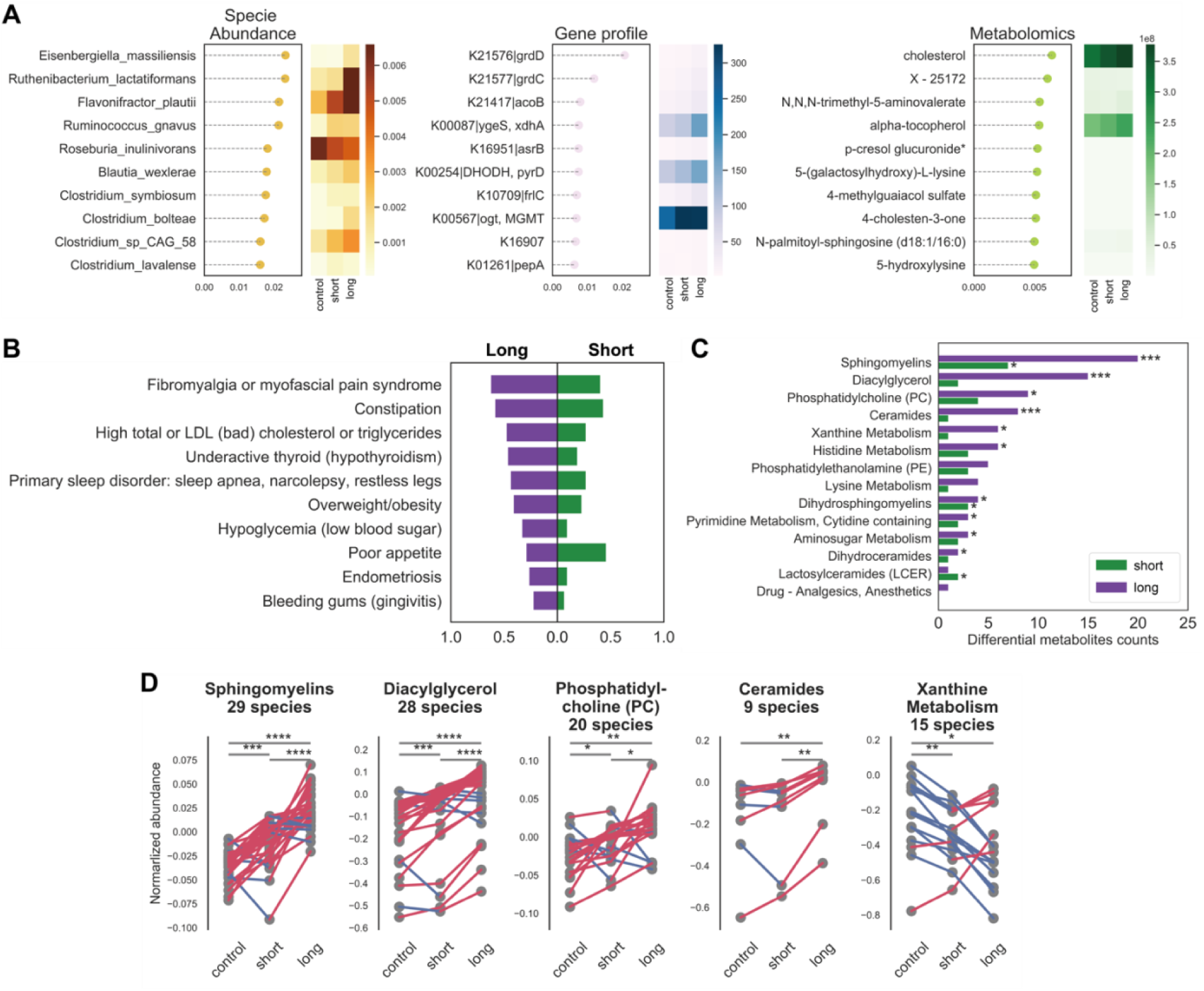
Phenotypical and metabolic abnormality are most pronounced in long-term patients. A) Multi gradient boosting models identified most important species, genes and metabolites differentiating controls, short-, or long-term ME/CFS. In each model, the top ten features are ranked by their contribution to the model on the y-axis, and the x-axis indicates the feature importance value. The heatmap shows the average feature abundance or relative abundance in each group. For full classification model performance (AUCs), see Figure S6. B) Naïve Bayesian model based on medical history records classified the stage of disease and identified nine significant clinical phenotypes in the long-term cohort and one significant phenotype in the short-term cohort. For each feature, the probability of experiencing the symptom in the long-term patients was presented to the left on the x-axis and the probability in the short-term patients was presented to the right. C) Overrepresentation analysis (ORA) on the plasma metabolome identified the most differential metabolites and pathways in the long-term group. For each pathway, two comparisons were conducted, control vs. short-term and control vs. long-term. P-values were computed by linear global t-test and the counts of differential metabolites presented on the x-axis. D) The trend of gradually changing metabolic irregularities along with the progression of disease are indicated by the difference between control, short-term and long-term cohorts. Here, each point represents the average normalized abundance for a given metabolite in the top five most abundant superfamilies, connected with a line that is colored by if increasing (red) or decreasing (blue). p-values were computed by Wilcoxon signed-rank test. p-value annotation legend: *: 0.01 < p <= 0.05, **: 0.001 < p <= 0.01, ***: 1e-04 < p <= 0.001, ****: p <= 1e-4

ORA with the metabolomics profiles identified the most striking differences between short- and long-term patients and controls. Unlike trends observed in the microbiome data with the greatest dysbiosis observed in short-term patients, long-term patients had more metabolites differentiating them from healthy controls, especially in sphingolipids and diacylglycerol metabolites (Figure 5C), confirming previous metabolomic findings. Interestingly, most metabolic species either decreased across experimental groups (control>short-term>long-term, e.g., xanthine;), or increased (control<short-term<long-term; e.g. sphingomyelins, diacylglycerol, phosphatidylcholine, and ceramides, Wilcoxon signed-rank test, p < 0.01), suggesting that metabolic irregularities associated with ME/CFS may gradually worsen over time (Figure 5D). Taken together, we postulate that long-term ME/CFS patients have developed a unique but stable pathophysiology characterized by more severe clinical symptoms and an array of altered host metabolic reactions.

### Gut and plasma butyrate is reduced in early-stage disease and is associated with host abnormal physiology

We noted in our metagenomics classification model a particular refrain of a depletion of butyrate-synthesizing microbes, including *Roseburia* and *F. prausnitzii* in ME/CFS. Butyrate is a major energy source for colonic epithelial cells and one of the main intestinal anti-inflammatory metabolites (^77,52^). It has been widely implicated in chronic disorders, including Crohn’s disease (CD) and ulcerative colitis (UC) (^53^). We thus performed a focused metagenomic and metabolomic analysis of the butyrate pathway to better understand its potential role in the crosstalk between the gut microbiota and host physiology in ME/CFS.

Strikingly, we found a depletion in plasma isobutyrate in the short-term group (Figure 6A, Wilcoxon rank-sum test, p = 0.03). Plasma levels of butyrate are directly linked to butyrate-producing gut microbiota because isobutyrate is absorbed primarily from the colon and translocated to the blood (^78^). Thus, we sought to link plasma metabolite abundance to gene-coding potential of the microbiome. We performed differential abundance analysis of KEGG pathways encoding butanoate synthesis (KEGG map00650) between patients and controls and found a clear difference in nearly every component of the pathway between patients and controls (Wilcoxon rank-sum test, p < 0.05, Figure 6C). Finally, we inferred gut butyrate abundance from metagenomic data using a Markov Chain Monte Carlo (MCMC) metabolite prediction algorithm (see Methods), in the absence of matched gut metabolomic data. Inferred concentrations of isobutyrate were significantly decreased in ME/CFS (Wilcoxon rank-sum test, p = 0.05), especially in the short-term group (Figure 6B, p < 0.01).

**Figure 6.**
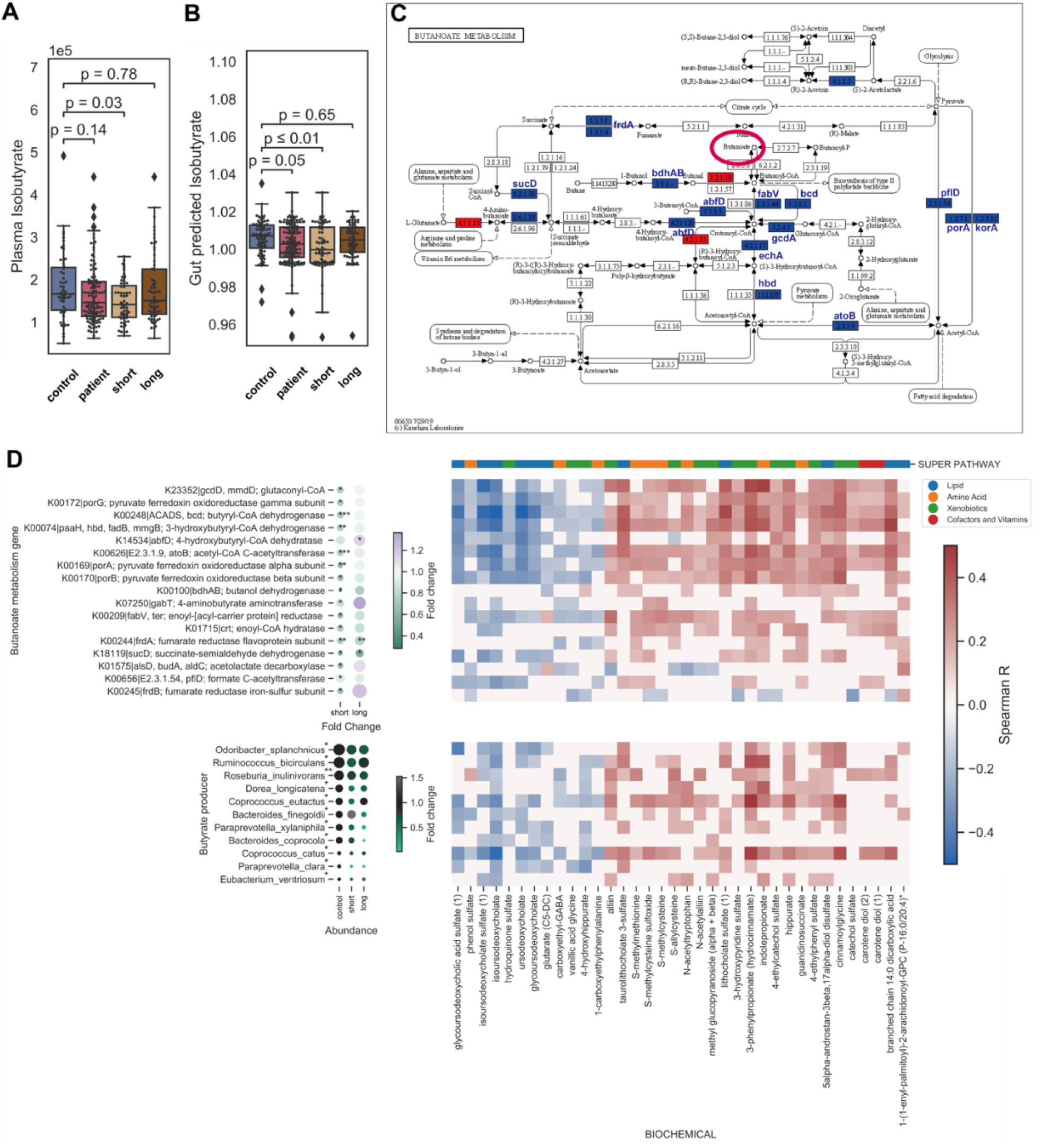
Limited microbial butyrate biosynthesis capacity associates with reduced plasma isobutyrate and multiple blood metabolites. In the blood and gut environment, decreased butyrate abundance in the short-term patient was indicated by: A) significantly reduced plasma isobutyrate normalized abundance; B) decreased predicted gut isobutyrate in patients, especially in short-term patients. Boxes show median relative abundance and interquartile ranges (IQR); whiskers specify ±1.5*IQR from the box’s quartile. P-values were computed by Wilcoxon rank-sum test. C) The reduced abundance of most key enzymes in the butanoate mechanism (KEGG pathway map00650) indicated a more limited microbial butyrate biosynthesis capacity in ME/CFS. Differentiating enzymes were colored and annotated on the map (decreased in blue and increased in red). D) Correlation of plasma metabolite normalized abundance and relative abundance of microbial butyrate biosynthesis features, with fold changes. Heatmap shows significant correlations (Spearman, p < 0.05) with the top bar indicating the metabolite superfamily. The top half shows the key enzymes in the KEGG butanoate pathway. On the left, different fold changes between the two patient cohorts (short-term vs. control and long-term vs. control, respectively) indicated a significant decrease in butyrate biosynthetic capacity in the early stages of ME/CFS. P-values were calculated in each group with Wilcoxon rank-sum test. Finally, the bottom half shows the correlation between the relative abundance of predicted butyrate producers and plasma metabolites. Microbes were ordered by relative abundance. For each microbe, the size of the dot indicates the mean abundance in each group and the color indicated fold change over the control group. P-value was computed by Kruskal–Wallis H test. p-value annotation legend: *: 0.01 < p <= 0.05, **: 0.001 < p <= 0.01, ***: 1e-04 < p <= 0.001.

Focusing on the enzymatic pathway (Figure 6C, 6D), in ME/CFS patients, particularly in the short-term group, there was a reduced abundance of most enzymes (*fabV, abfD, bcd, gcdA*, and *echA*) that produced crotonyl-CoA, an essential intermediate in the fermentation of butyric acid(^79^). The reduction of crotonyl-CoA might also alter the metabolism of lysine and tryptophan, as it is also reported to be necessary for metabolism of fatty acids and amino acids(^80^). The reduced relative abundance of pyruvate ferredoxin oxidoreductase (*porA* and *korA*) and acetyl-CoA C-acetyltransferase (*atoB* and *pflD*) could hint at a deficit in production of acetyl-CoA, which participates in multiple essential metabolic mechanisms, including the acetate and propionate biosynthesis pathways(^81^). Similarly, the reduced levels of both succinate dehydrogenase (*frdA)* and fumarate reductase (*sucD*) observed could result in dysbiosis of succinate, an essential intermediate in the synthesis of propionate by gut bacteria and an abundant product of microbial fermentation of dietary fiber(^82^).

Finally, we performed a correlation analysis with the relative abundance of 1) butyrate producing microbes or 2) KEGG enzymes with plasma metabolites to identify the degree to which this pathway may influence circulating metabolite levels (Figure 6D, Table S8, see Methods). From the thousands of plasma metabolites tested, we identified 24 positive correlations, including moieties from propionates, succinates, tryptophans, and hippurates, consistent with results of our differential abundance analysis, as well as 12 negative correlations, including sulfate and ursodeoxycholate moieties. The former suggests that changes in the ability of the gut microbiome to metabolize or synthesize short-chain-fatty acids (SFCA) is reflected in dysbiosis of these and related metabolites in plasma. In addition, hippuric acid, produced by gut bacterial metabolism of dietary components, has been previously associated with gut microbial community diversity and positively correlated with butyrate producers *Clostridiales* sp. and *F. prausnitzii* (^83^). For negatively associated metabolites, both of them are microbiota-derived metabolites in the plasma (^84^). Ursodeoxycholate metabolites, one secondary bile acid, was a side product from lipid absorption process, which conjectured a possible link between the gut microbiome and dysbiosis observed in lipid-related metabolites(^85^).

### The interactions between gut microbes and host plasma metabolites in ME/CFS

Finally, to identify additional links between the gut microbiome and the plasma metabolome, we took a step back and performed several global co-association analyses with plasma metabolites and microbiome features. First, we performed an association analysis with the most abundant gut bacteria identified: three SCFA producers - *F. prausnitzii*, an important butyrate producer identified as a biomarker in our classification models, *A. putredinis*, and *E. coli*, which has significant immunomodulatory and metabolic impacts(^86,87^) – as having the greatest number of significant correlations with plasma metabolites, conceptually consistent with our results thus far (Figure 7A).

**Figure 7.**
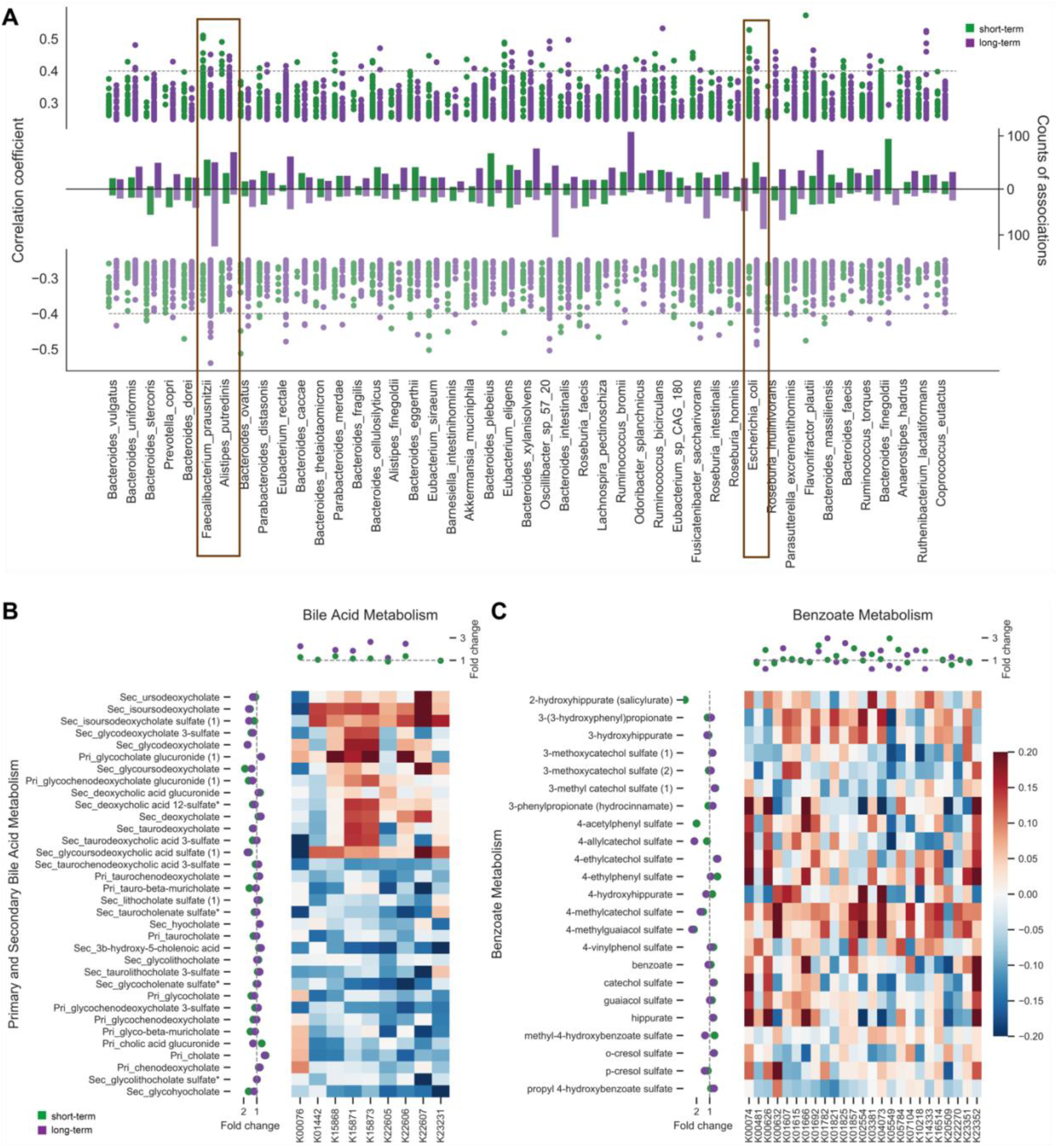
The multi–‘omics association structure of the host metabolome and gut microbiome in the short-term and long-term cohorts. A) Gut microbes were differentially associated with the plasma metabolome in short- and long-term cohorts. Global association studies (Spearman correlation) were applied between every gut microbe and every plasma metabolite in the two patient cohorts to capture interaction patterns in the different stages of the disease. The most abundant species were ordered by their relative abundance on the x-axis with their significant associations (p < 0.05) in both groups (short-term in green, long-term in purple) summarized in the y-axis. For each bacteria, the positive and negative associations are displayed in two directions of the y-axis (top and bottom, respectively) with their coefficient (R^2^) as dots, and the total count of statistically significant associations is shown as the bar in the middle. For example, *Faecalibacterium prausnitzii, Alistipes putredinis*, and *Escherichia coli* have significantly different numbers of correlated metabolites and are highlighted in frame. B) The correlation pattern between normalized abundance of plasma bile acid metabolism and relative abundance of the microbial secondary bile acid biosynthesis pathway (Spearman) with fold changes are shown. The x-axis includes key enzymes from the microbial bile acid pathway (KEGG map00121) with their fold changes in the two patient cohorts annotated on the top. The y-axis indicates bile acids in the plasma with fold changes on the left. The heatmap shows that most secondary bile acids were positively correlated with the microbial bile acid pathway, while most primary bile acids were negatively associated. C) Fold changes as in B) are shown for the positive correlation between plasma benzoate metabolism and microbial benzoate pathway (Spearman). The x-axis includes key enzymes from the microbial benzoate pathway (KEGG map00362) with their fold changes annotated on the top. The y-axis indicates the bile acids in the plasma with fold changes on the left.

We then associated overall community diversity with metabolomic sub-pathways, as plasma metabolites have been shown to predict gut microbial diversity with implications for aging. We found six host metabolic pathways associated with the gut microbiome (Figure S7). Two positive correlated examples would be sphingomyelin (Spearman, R = 0.16, p < 0.05) and vitamin A metabolism (Spearman, R = 0.26, p < 0.01). Both sphingolipid metabolites and vitamin A shape host-microbiome interactions via the immune system, either by modulating trafficking and function of immune cells (^88^) or mediating influencing immune homeostasis in the gut. (^89^). Conversely, diversity was also correlated with the regulation of primary/secondary bile acids (Figure S7, Spearman, R = -0.18 & -0.15, p < 0.05). Primary bile acids are small molecules synthesized by the liver and translocated to the colon for lipid digestion. Though 95% of primary bile acids are reabsorbed by the gut, some *Clostridales sp*. can convert primary bile acids to secondary bile acids in the colon (^90^). With this negative correlation with primary bile acids, and a positive correlation in the secondary bile acids and overrepresentation of the bacterial secondary bile acid biosynthesis pathway (KEGG map00121, Figure 7B), we conjecture that secondary bile acids, synthesized by the gut microbes from primary bile acids, is upregulated in ME/CFS. Another microbial metabolic pathway known to impact host metabolism, benzoate metabolism, was also positively associated with plasma metabolites (Figure S7, Spearman, R = 0.24, p = 0.01). Benzoic acid is synthesized through fermentation by colonic microbiota of dietary aromatic compounds and is then conjugated with glycine in the liver to produce Hippurate(^91^), which is associated with the reduction of butyrate as discussed above. The positive correlation that we observed between plasma benzoate and the Simpson diversity, as well as with the gene abundances of the bacterial benzoate degradation pathway (KEGG map00362, Figure 7C) also hinted at a reduced benzoic intermediate and hippurate. This interaction has also been reported in type 2 diabetes and CD (^92,93,94^).

In short, decreased gut microbiome diversity was, in several cases, linked with changes in the gene-coding potential of key bacterial functions associated with host physiological and metabolic alterations. Collectively, these data suggest new microbial links to changes in the homeostasis of circulating metabolites that can have widespread impacts on immunity, metabolism, and other functions that are dysbiotic in ME/CFS.

## Discussion

Here, we performed the first large-scale multi-‘omics investigation integrating detailed clinical and lifestyle data, gut metagenomics, and plasma metabolomics in short- and long-term ME/CFS patients compared to healthy controls. Several studies have reported a disrupted gut microbiome in ME/CFS (^11,13^) as well as changes in blood cytokine and metabolite levels. However, our cohort design differentiating short- and long-term patients was important to identify microbial and metabolic features that may contribute to disease progression.

Notably, the most significant microbial dysbiosis occurred in short-term ME/CFS. Compositional differences in short-term ME/CFS were consistent with previous studies using lower resolution sequencing (^95^), identifying a broad reduction in microbial diversity (^96^), alterations in the ratio of Bacteroides:Firmicutes microbiota (^10^), and increased heterogeneity of low abundance organisms. This latter feature has recently been associated with the microbiome of frail older adults and supports its general association with reduced health outcomes(^39^). There are several potential explanations for this early-stage dysbiosis. First, short-term patients suffer more GI disturbances, and gut microbiome changes may reflect these environmental changes. Second, it is possible that patients may try a range of interventions that impact their gut microbiome, which is dynamic and influenced by numerous intrinsic and extrinsic factors, including age and diet(^97^). We examined age as a covariate, and consistent with our previous study, found that age can influence gut microbiome composition. Diet can not only dramatically shift gut microbial community composition, but it can alter the metabolic potential of the microbes and production of immunomodulatory metabolites. We also analyzed the detailed dietary metadata, showing that most dietary habits are comparable among our cohorts, except infrequent sugar intake in our short-term patients, which might also contribute to the microbial differences observed in early stages of disease (Table S1, Figure S1).

The return of the gut microbiome of long-term patients to a configuration more similar to healthy controls (with notable differences nonetheless, in low abundance species and in heterogeneity) as well as the reduced occurrence of gastrointestinal illness in this cohort, suggests a return to a relative homeostasis. However, we conjecture that microbial dysbiosis seen in short-term patients may have cumulative and long-term effects, where damage may be caused by an initial trigger, resulting in cascading events. Long-term patients, despite their relatively more ‘control-like’ gut microbiome, have more severe clinical symptoms and metabolic dysbiosis. Thus, we hypothesize that ME/CFS progression may begin with loss of beneficial microbes, particularly SCFA producers, resulting in more pervasive gastrointestinal phenotypes that is later reflected in plasma metabolite levels. Individual-specific changes then lead to the irreversible metabolic and phenotypic changes and unrecoverable ME/CFS.

The abnormalities in the short-term cohort could result in potential increases in aberrant translocation of microbial metabolites that could affect host immune and metabolic processes. For example, one of the changes we noted in short-term patients was a reduction in potential immunomodulatory organisms (butyrate and tryptophan producers, e.g.. *Faecalibacterium prausnitzii*), which we speculate could lead to long-term metabolic dysbiosis. The reduced prevalence of the butanoate synthesis pathway and the reduced relative abundance of butyrate-producing bacteria among all patients, but especially in the short-term group, suggested a loss of butyrate in the intestinal environment. This was consistent with the decrease of isobutyrate measured in the blood. We note however, that our long-term cohort is slightly but significantly older than the short-term cohort (though age-matched in our healthy controls), which could contribute to some of the phenotypes, particularly clinical abnormalities, observed.

Butyrate, tryptophan, and other microbial metabolites have been linked to mucosal immune regulation, and our team previously showed a striking immune dysbiosis in different blood immune markers, including changes in the functional capacity of mucosal associated invariant T (MAIT) cells and Th17 cells, and a decrease in the frequency of CD8+ T cells and natural killer cells in long-term ME/CFS patients (^44^). This is an exciting association because each of these cell types have been linked to bacterial or fungal infections, respond to microbial metabolites, and have been linked to the pathogenesis of autoimmune or chronic inflammatory diseases. Thus, it is possible that the microbiome primes or sustains an aberrant immune response following disease onset. This is supported by an observed shift from a predominantly Th1 to Th2 immune response in ME/CFS (^98^).

There is currently no standard diagnostic test for ME/CFS because of many phenotypes of ME/CFS are shared with other disorders(^5,6^), such as fibromyalgia. Here, our integration of multiple ‘omics data significantly increased classification accuracy and identified microbial and metabolic features that could pinpoint potential hypotheses for further investigation and therapeutic strategies. Longitudinal sampling of short-term patients, particularly as they progress to long-term disease, would help to untangle directionality of microbial dysbiosis and potential effects on the blood metabolome. For example, recent studies performing large-scale associations with the gut microbiome and blood metabolome have identified that a significant fraction (upwards of 15%) of blood metabolites can be predicted by gut microbiome composition(^99^). However, understanding of the temporal nature of this association is limited. Here, our ‘omics workflows could be one of the guiding frameworks to intergrading microbiome, metabolome, and host phenotypes, and thus, bring a more comprehensive understanding to host-microbiome interactions.

Finally, we believe that recent potential associations between the chronic immune dysfunctions in ME/CFS patients and ‘long COVID’ increase the relevance of the results reported here. ‘Long COVID’ refers to phenotypes suffered by numerous patients infected by SARS-CoV-2 (COVID-19) that have ‘recovered’, but did not return to full health. Notably, it manifests as numerous phenotypical abnormalities shared with ME/CFS, including lingering chronic fatigue and myalgias. In ME/CFS, key symptoms might also be triggered by acute infections including SARS coronavirus, MERS (^100^), the Epstein-Barr Virus (EBV) (^101^), or other agents, and such infections were reported to be more frequent in the medical histories of our patient cohort, even preceding the onset of ME/CFS symptoms (^102,103^) Understanding the biological mechanisms underlying ME/CFS may now have further urgency and generalizability in the worldwide COVID-19 pandemic. Taken together, we have established a framework to study host-microbiome interactions leveraging ‘omics to identify host and microbial metabolites and functions implicit in ME/CFS, presenting a rich clinical and ‘omics dataset to further mechanistic hypotheses to better understand this debilitating disease.

## Methods

### Cohort and study design

All subjects were recruited at Bateman Horne Center, Salt Lake City, UT, based on who met the 1994 CDC Fukuda (Fukuda et al., 1994) and/or Canadian consensus criteria for ME/CFS (Carruthers, 2007). Healthy controls were frequency-matched to cases on age, sex, race/ethnicity, geographic/clinical site, and season of sampling. Patients or controls taking antibiotics or who had any infections in the prior one month, or who were taking any immunomodulatory medications were excluded from the study. The study was approved by The Jackson Laboratory IRB (Study number 17-JGM-13) and written informed consent and verbal assent when appropriate were obtained from all participants in this study. We enrolled a total of 149 ME/CFS patients (of which 74 had been diagnosed with ME/CFS <4 years before recruitment and 75 had been diagnosed with ME/CFS >10 years before recruitment) and 79 healthy controls. Subject characteristics are shown in Supplemental Table 1.

### Participants

149 ME/CFS patients who had been seen at Bateman Horne Center (Salt Lake City, UT) for routine clinical care between February 2018 and September 2019 and 79 matched HCs were recruited for the study. The 149 MECFS subjects included 74 sick with ME/CFS for <4 years and 75 sick for greater than 10 years. The age range of ME/CFS participants and HCs was 18-65 years at the time of informed consent. HCs were matched with <4 ME/CFS participants by age (± 5 years), gender and ethnicity. Enrolled ME/CFS participants were required to fulfill the International Chronic Fatigue Syndrome Study Group research criteria(^104^), the Canadian Consensus Criteria(^105^), and the IOM clinical diagnostic criteria(^106^). HCs were recruited from the Salt Lake City metropolitan area using advertisements posted on social media, the clinic webpage or by phone contact with a volunteer pool from previous studies. HCs were considered generally healthy and between 18 to 65 years of age. HCs were excluded if they fulfilled ME/CFS diagnostic criteria or had a history of illness, had a BMI>40 or had been treated with long-term (longer than 2 weeks) antiviral medication or immune modulatory medications within the past 6 months or had been treated with short-term (less than 2 weeks) antiviral or antibiotic medication within the past 30 days.

### Clinical metadata collection and preprocess

Clinical symptoms and baseline health status was assessed on the day of physical examination and biological sample collection from both case and control subjects. For each participant, we collected demographic information (including age, gender, diet, race, family, work, and education), medical histories, and three questionnaires regarding the general physical and mental health condition (RAND-36 form), sleep quality (PSQI form) and gastrointestinal health (GSRS form). The summary of analyzed clinical features and questionnaires are shown in supplemental Table 2. Age and diet were analyzed and discussed as potential confounders (Figure S1 and S4). Medical histories were simplified into binary features (0 - no records, 1, had/having the disease) and further constructed naïve Bayesian classification models. Every questionnaire was transformed into a 0–100 scale to facilitate combination and comparison wherein a score of 100 is equivalent to maximum disability or severity and a score of zero is equivalent to no disability or disturbance.

### Plasma sample collection and preparation

Healthy and patient blood samples were obtained from Bateman Horne Center, Salt Lake City, UT and approved by JAX IRB. One 4 mL lavender top tube (K2EDTA) was collected, and tube slowly inverted 8-10 times immediately after collection. Blood was centrifuged within 30 minutes of collection at 1000 x g with low brake for 10 minutes. 250 uL of plasma was transferred into three 1 mL cryovial tubes, and tubes were frozen upright at -80°C. Frozen plasma samples were batch shipped overnight on dry ice to The Jackson Laboratory, Farmington, CT, and stored at -80°C.

### Plasma untargeted metabolome by UPLC-MS/MS

Plasma samples were sent to Metabolon platform and processed by Ultrahigh Performance Liquid Chromatography-Tandem Mass Spectroscopy (UPLC-MS/MS) following the CFS cohort pipeline. In brief, samples were prepared using the automated MicroLab STAR® system from Hamilton Company. The extract was divided into five fractions: two for analysis by two separate reverse phases (RP)/UPLC-MS/MS methods with positive ion mode electrospray ionization (ESI), one for analysis by RP/UPLC-MS/MS with negative ion mode ESI, one for analysis by HILIC/UPLC-MS/MS with negative ion mode ESI, and one sample was reserved for backup. QA/QC were analyzed with several types of controls were analyzed including a pooled matrix sample generated by taking a small volume of each experimental sample (or alternatively, use of a pool of well-characterized human plasma), extracted water samples, and a cocktail of QC standards that were carefully chosen not to interfere with the measurement of endogenous compounds were spiked into every analyzed sample, allowed instrument performance monitoring, and aided chromatographic alignment. Compounds were identified by comparison to Metabolon library entries of purified standards or recurrent unknown entities. The output raw data included the annotations and the value of peaks quantified using area-under-the-curve for metabolites.

### Metabolome enrichment study

From the raw data (peaks area-under-the-curve), we first kept metabolic features present in >50% of the samples for further analysis, and missing values were imputed using the k-nearest neighbor method. We then normalize the raw area counts by rescaling the median of each metabolite to be 1. We first applied qualitative enrichment analysis (Over Representation Analysis ORA) in two patient cohorts (short-term vs. control, and long-term vs. control). We conducted Wilcoxon rank-sum test on all metabolites and counted the significantly differential metabolites in every sub pathway. Fisher test with Benjamini-Hochberg adjustment was followed to identify the distributions of the over-representated genes in the pathway. For every sub pathway, we also applied a global quantitative enrichment analysis with linear *globaltest* in R to compute the association between a group of metabolites from that pathway and the duration of disease (control, short-term, and long-term).

### Fecal sample collection and DNA extraction

Stool was self-collected at home by volunteers using a BioCollector fecal collection kit (The BioCollective, Denver, CO) according to manufacturer instructions. The study participants also added a portion of the stool sample to an OMNIgene•GUT tube (DNA Genotek, OMR-200) following manufacturer instructions for preservation for sequencing prior to sending the sample in a provided Styrofoam container with a cold pack. Upon receipt, stool and OMNIgene samples were immediately aliquoted and frozen at –80°C for storage. Prior to aliquoting, OMNIgene stool samples were homogenized by vortexing (using the metal bead inside the OMNIgene tube), then divided into 2 microfuge tubes, one with 100μL aliquot and one with 1mL. DNA was extracted using the Qiagen (Germantown, MD, USA) QIAamp 96 DNA QIAcube HT Kit with the following modifications: enzymatic digestion with 50μg of lysozyme (Sigma, St. Louis, MO, USA) and 5U each of lysostaphin and mutanolysin (Sigma) for 30 min at 37 °C followed by bead-beating with 50 μg 0.1 mm of zirconium beads for 6 min on the Tissuelyzer II (Qiagen) prior to loading onto the Qiacube HT. DNA concentration was measured using the Qubit high sensitivity dsDNA kit (Invitrogen, Carlsbad, CA, USA).

### Metagenomic shotgun sequencing

Illumina libraries were created using Nextera XT DNA Library Prep Kit (Illumina, San Diego, CA, USA) with reduced reaction volumes: 200pg of DNA were used (160 pg/μL × 1.25 μL), and tagmentation and PCR reagent volumes were reduced to 1/4 of the standard volumes. Tagmentation and PCR reactions were carried out according to the manufacturer’s instructions. The reaction mixtures were then adjusted to 50 μL by adding dH2O, and the AMPure (Beckman Coulter) Cleanup was carried out as per the manufacturer’s instructions. Libraries were then sequenced with 2 × 150 bp paired end reads on Illumina HiSeq2500 and NovaSeq6000. For quality control, we used standard mock sample sequenced by HiSeq and NovaSeq to predict the taxonomy composition and confirmed that there was no difference between these two sequence techniques after the normalization. Sequencing adapters and low-quality bases were removed from the sequencing reads using scythe (v0.994) and sickle (v1.33), respectively, with default parameters. Host reads were then filtered by mapping all sequencing reads to the hg19 human reference genome using bowtie2 (v2.2.8), under “very-sensitive” mode. Unmapped reads were used for downstream analyses.

### Taxonomic and KEGG gene profiling of metagenomics samples

Taxonomic compositions were profiled using Metaphlan3.0 (^107^) and the species whose average relative abundance > 1e-5 were kept for further analysis, giving 384 species. The gene profiling was computed with USEARCH(v8.0.15)(^108^) (with parameters: evalue 1e-9, accel 0.5, top_hits_only) to KEGG Orthology (KO) database v54, giving a total of 9452 annotated KEGG genes. The reads count profile was normalized by DeSeq2 in R(^109^).

### Microbial community structure analysis

The overall community structures was examined by Correspondence Analysis (PCoA) and PERMANOVA were performed in R with the *adonis* function in the R package vegan to analyze the partitioning of variation giving potential confounders including age and gender. The heterogeneity index (Inter-individual divergence) and community indexes including chao1, evenness(evar), rarity(low_abundance) and inequality(dominance_gini) were computed by R package microbiome. The species replication rate was predicted using GRiD with default settings(^48^).

### Gut metabolic status prediction

Here, we adapted from the MAMBO(^110^) pipeline and predicted 224 metabolites status of the microbial community in our cohort. We first constrained the Genome-scale metabolic models (GSMMs) community of all 384 microbes identified in the species profile. We started with the same gut metabolic environment for all samples giving randomized 224 metabolites initialization and using a Markov chain Monte Carlo (MCMC) to sample metabolites. In each step, we sampled one metabolite and used the Flux balance analysis to model reaction flux with a slight change of the target metabolites and accepted the step only if the probed growth rate is correlated with the species relative abundance. Samples were first subjected to 100,000 search steps, and 100,000 steps were subsequently added until a high Pearson correlation (ρ 0.6) with the target metagenomic abundance profile was achieved. Finally, the 10% time points with the highest Pearson correlation scores between the biomass profile and the metagenomic abundance profile were averaged, yielding a robustly predicted metabolome.

### Multi-‘omics classification models

To identify phenotypical, metagenomic, and metabolomic markers of the onset(control/patient) of the disease, we constructed a naïve Bayesian classification model with medical history records and three individual classification models based on the species abundance, normalized KEGG gene abundance, and normalized metabolite profile and one combination multi-omics model with all top ten features collected from each model. We also tested four different classification methods, LASSO logistic regression, Support vector machine (SVM), Random Forest (RF), and Gradient Boosting (GDBT). The same Multi-‘omics classification model system was also applied to classify the duration(control/short-term/long-term) of the disease.

All analyses were carried out using the Python package ‘scikit-learn’. Normalized KEGG gene and normalized plasma metabolome were standardized (by centering to mean 0 and dividing by the standard deviation of each feature) before fitting into the models. The models were optimized by five-fold RandomizedSearchCV to probe the best parameters giving lists of candidates. Models were then validated by 10-fold stratified cross-validation testing (we resampled dataset partitions 10 times). In each test, the accuracy of the model was examined using ROC (area under the curve). The comparison among models showed that GDBT outcompeted the rest three and reached the best performance. The parameters for the optimized GDBT model: sklearn.ensemble.GradientBoostingClassifier(ccp_alpha=0.0, criterion=‘friedman_mse’, init=None, learning_rate=0.05, loss=‘deviance’, max_depth=7, max_features=2, max_leaf_nodes=None, min_impurity_decrease=0.0, min_impurity_split=None, min_samples_leaf=9, min_samples_split=20, min_weight_fraction_leaf=0.0, n_estimators=100, n_iter_no_change=None, presort=‘deprecated’, random_state=1015, subsample=0.8, tol=0.0001, validation_fraction=0.1, verbose=0, warm_start=False). In the Gradient boosting model, two steps were carried out. In the first step, the model was constructed using each of the three profiles (species abundance, KEGG gene abundance, and metabolite profile) individually to compute the feature importance as the feature contributions to the classification. In the second step, the collective model was constructed using a combination of top ten important features determined from the species relative abundance model + top ten important features determined from the KEGG gene model + top ten important features determined from the plasma metabolite model.

### Interaction study and targeted pathway analysis

Spearman correlations (ρ) of non-zero values were used for all correlation coefficients. To understand the interaction between host and microbe, we started with a global correlation analysis by pairwise correlations between species and plasma metabolites for each cohort (control, short-term, and long-term). The positive and negative significant correlations (p < 0.05) for the top 50 abundant species were summarized by cohort. We then conducted linear regression analysis with the gut microbiome community structure index (Simpson diversity) and the sub pathways in the plasma metabolome. The relative abundance of the metabolic pathway was indicated by the mean of all metabolites in the pathway. We identified several metabolic pathways that significantly related with the microbial community. Thus, the targeted pathway analysis was applied for the identified pathway as well as some important microbial metabolic metabolisms, like the butyrate pathway. We computed the Spearman correlation for the gut microbial KEGG pathway and their paired plasma metabolic pathway along with the fold change of the elements, genes, and metabolites, respectively.

### Statistical Analysis

The dimensionality reduction analysis was conducted by Principal Correspondence Analysis (PCoA) using sklearn.manifold.MDS function for both gut microbiome Bray-Curtis dissimilarity distance matrix and normalized plasma metabolome profile. The statistically significant differences among independent groups (healthy/patient/short-term/long-term) were determined by nonparametric test using Wilcoxon rank-sum test two-sided with Bonferroni correction. The average abundances of each species and metabolites were determined to be significantly elevated or depleted in short-term or long-term group by pairwise nonparametric comparison using Wilcoxon signed-rank test with Bonferroni correction. Chi-squared test was used to compare the infection frequencies between healthy and patient group. P value annotations: ns: p > 0.05, *: 0.01 < p <= 0.05, **: 0.001 < p <= 0.01, ***: 1e-04 < p <= 0.001, ****: p <= 1e-04.

## DATA AVAILABILITY

The dataset related to this article can be found at the Short Read Archive (SRA) under the BioProject accession number TBD.

## ACKNOWLEDGEMENTS AND FUNDING

We are thankful to the Oh, Unutmaz, and Li laboratories for inspiring discussions and acknowledge the contribution of the Genome Technologies Service at The Jackson Laboratory for expert assistance with sample sequencing for the work described in this publication. We also thank the clinical support team at the Bateman Horne Center and all the individuals who participated in this study. This work was funded by 1U54NS105539. JO is additionally supported by the NIH (1 R01 AR078634-01, DP2 GM126893-01, 1 U19 AI142733, 1 R21 AR075174).

## AUTHOR CONTRIBUTIONS

Conceptualization: DU, JO, SDV, LB, RX; Data Curation: RX, CG, SDV, LB; Formal Analysis: RX; Funding Acquisition: DU, JO, SDV, LB; Clinical sample design and collection: SDV, LB; Investigation: RX, CG, EF, SDV, LB; Project Administration: JO, DU, LB, SDV, CG; Resources: DU, JO, SDV, LB; Supervision: JO; Visualization and Writing: RX, JO; Writing - Review and Editing: RX, CG, SDV, LB, DU, JO.

