## Supplemental Figures for "Multi-‘omics of host-microbiome interactions in short- and long-term Myalgic Encephalomyelitis/Chronic Fatigue Syndrome (ME/CFS)"

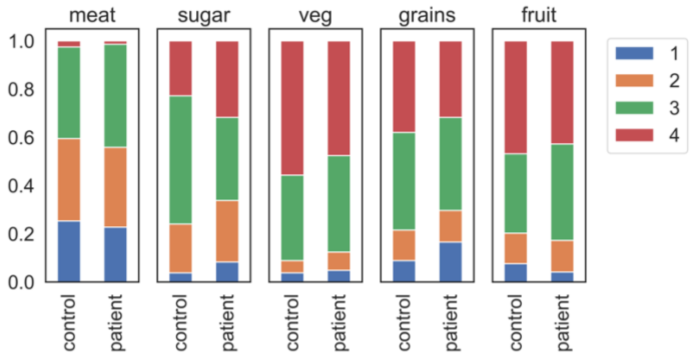


**Supplemental Figure 1. The dietary distributions of control and patient cohorts.** The dietary questionnaire was summarized into five categories and their frequency in the last week. Categories: Meat - red meat; sugar - desserts, sweets, soda, or juice; veg – fresh vegetables; grains, whole grains (e.g., oatmeal, quinoa, whole wheat products); fruit - fresh fruit. Frequency: 1, Never; 2, Once; 3, 2 to 5 times; 4, Daily.

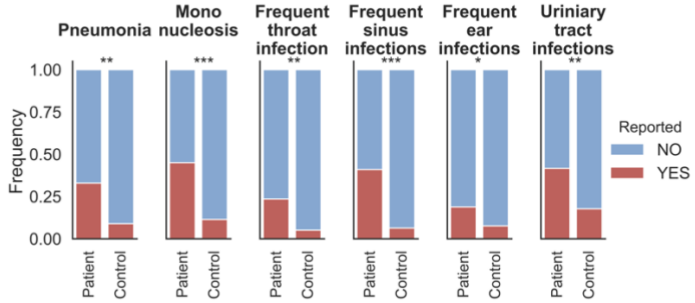


**Supplemental Figure 2. ME/CFS patients reported significantly more infection-related histories.** More than 30% ME/CFS patients reported pneumonia, mononucleosis, frequent sinus infections and urinary tract history. The p value was computed by Chi-squared test. p-values annotations: *: 0.01 < p <= 0.05, **: 0.001 < p <= 0.01, ***: 1e-04 < p <= 0.001

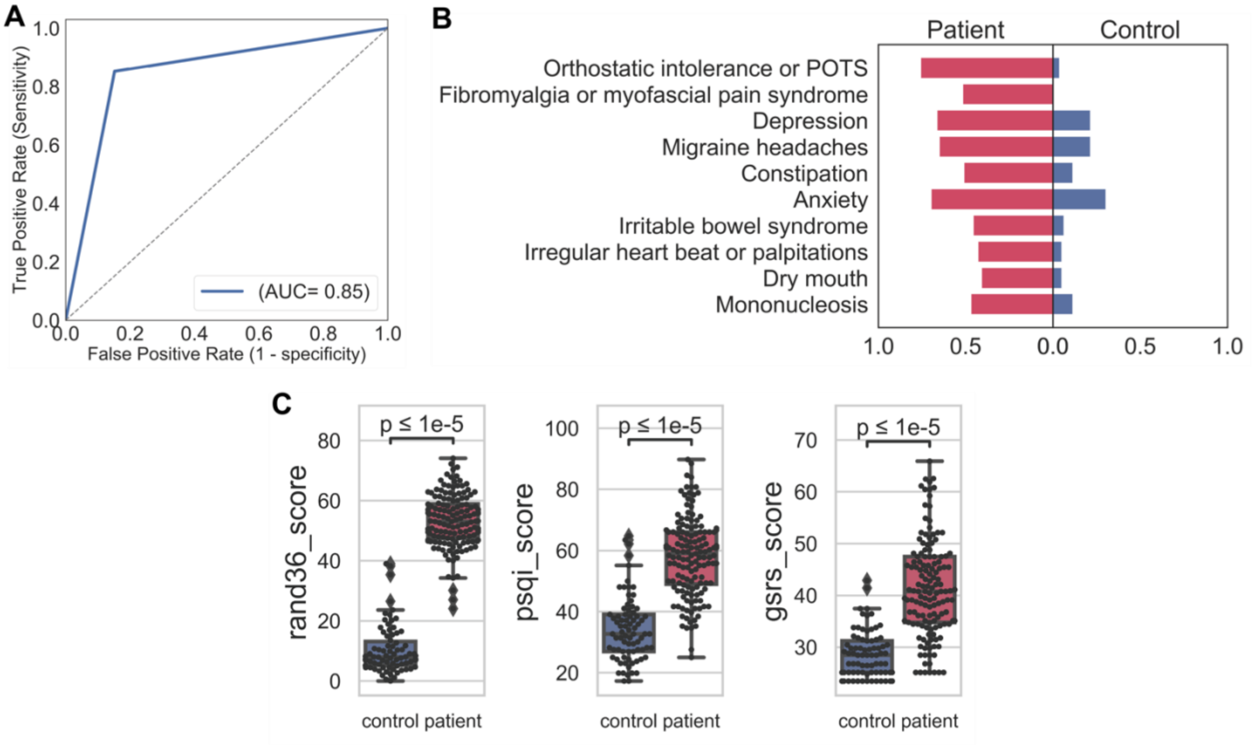


**Supplemental Figure 3. The host phenotype in ME/CFS.** A) The performance of the naïve Bayesian classification model based on medical history records to identify clinical features that discriminate healthy controls vs. patients. The area under the curve, AUC = 0.85. B) Top ten predicted features and their probabilities to discriminate both cohorts were presented in separate directions on the x-axis. For each feature, the probability of experiencing the symptom in the patients was presented to the left on the x-axis and the probability in the controls was presented to the right. C) Based on our scoring system, patients had significantly more anomalous mental and physical health conditions identified by higher rand36, Pittsburgh sleep quality, and gastrointestinal symptom rating scale scores (Table S2). p-values were computed by Wilcoxon rank-sum test.

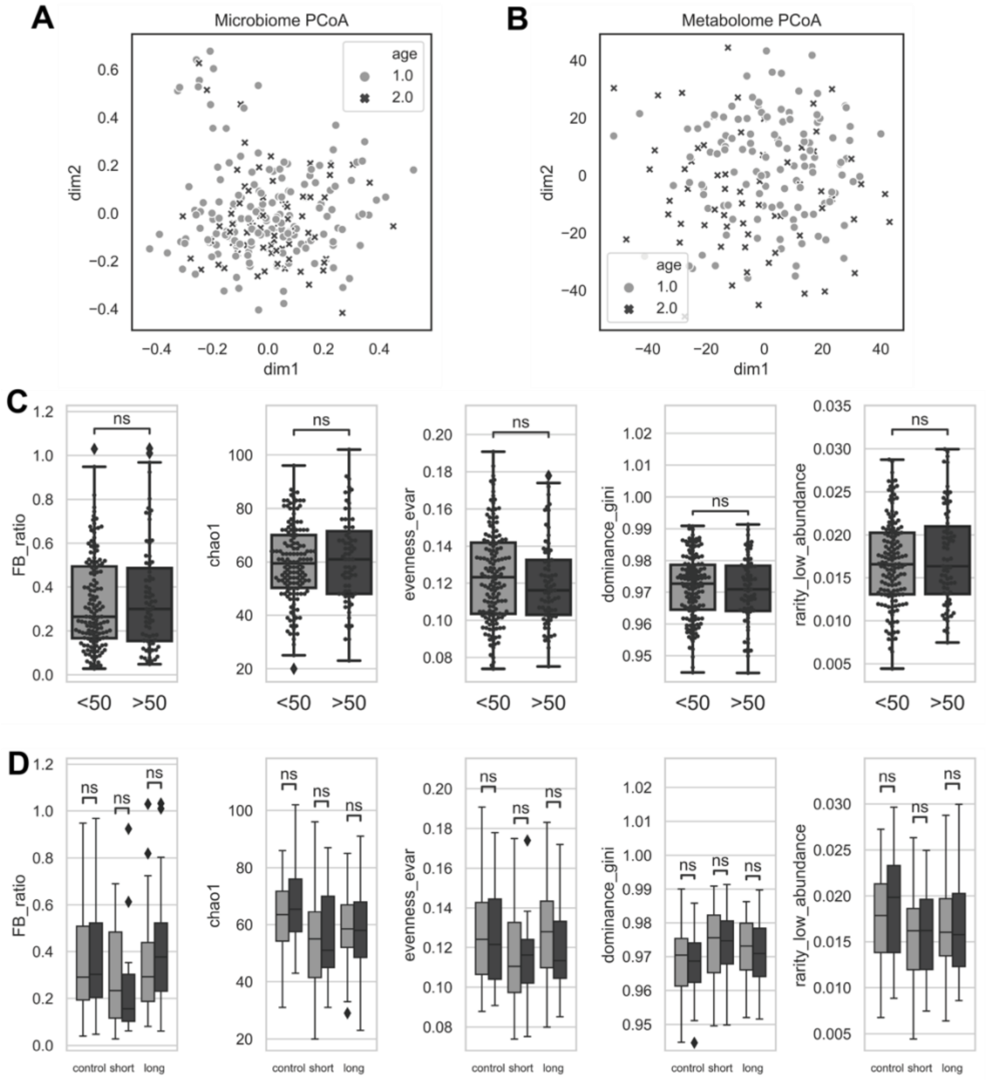


**Supplemental Figure 4. Age is not a significant confounder of the gut microbiome and plasma metabolome.** A) The Principal Correspondence Analysis (PCoA) based on gut microbiome-derived Bray-Curtis dissimilarity distance. The contribution of age was not significant with a p-value > 0.05 (PERMANOVA, see Methods). B) Principal Correspondence Analysis (PCoA) based on normalized plasma metabolome profile. In C) and D), microbial community structure was not significantly different between young (<50 years old) and old (>=50 years old) in C) all cohorts and D) cohort by disease stage (control, short-term, and long-term), respectively. Community structure was indicated by FB_ratio (Firmicutes:Bacteroides ratio), Chao 1 index (richness), evenness_evar (Smith and Wilson's Evar index), dominance_gini (Gini index of the dominant species >0.2% relative abundance), and rarity_low_abundance (proportion of the least abundant species <0.2% relative abundance). p-values were computed by Wilcoxon rank-sum test. ns, not significant.

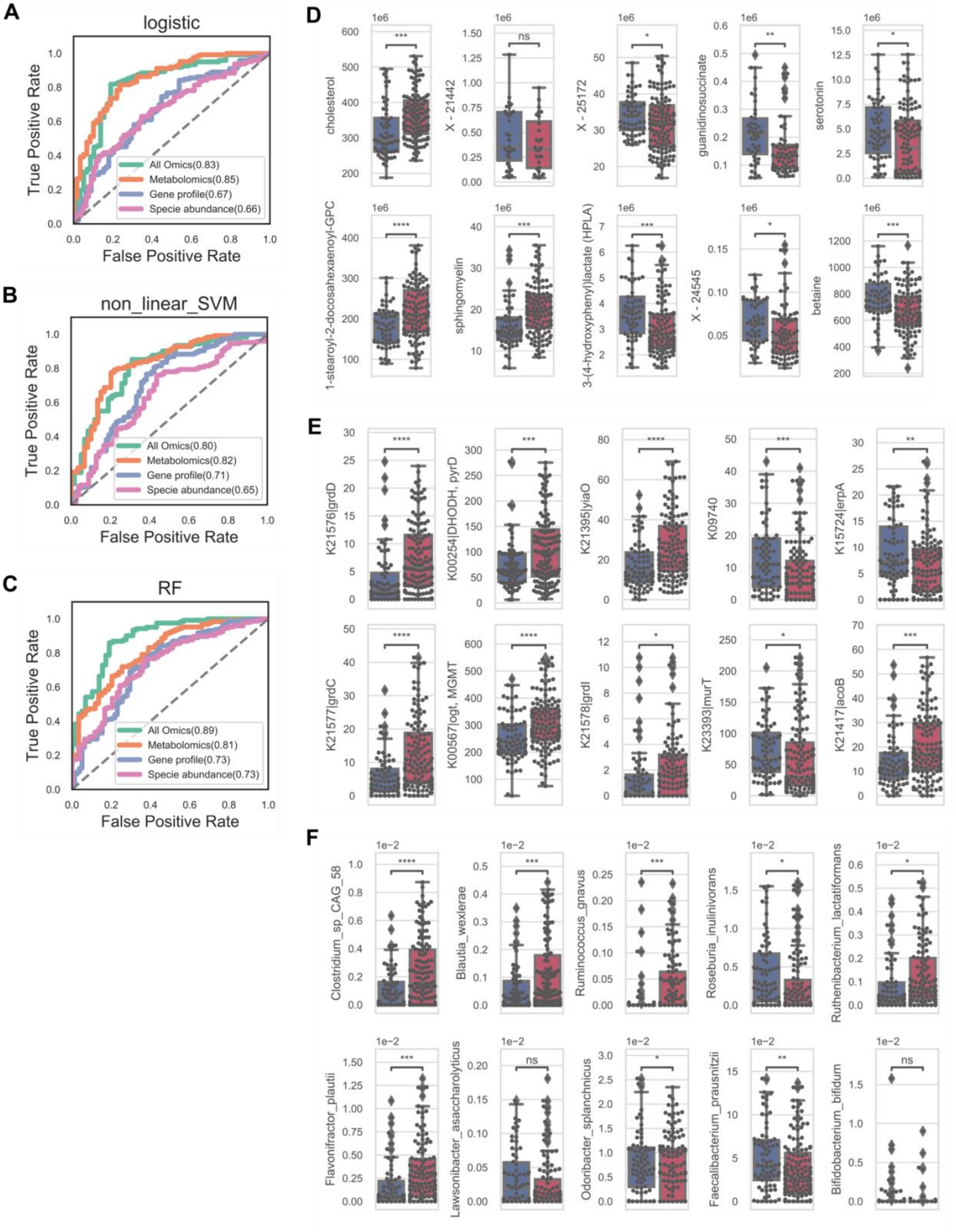


**Supplemental Figure 5. Multi-'omics models to classify the onset of ME/CFS (control vs. patient).** A-C) Performance of the classifiers using area under the curve (AUC) was evaluated using 10 randomized and 10-fold cross-validations for each model: LASSO logistic regression, Support vector machine (SVM) and random-forest (RF) models. Models were designed based on species relative abundance (pink), KEGG gene profile (blue) or plasma metabolome (orange) individually, or a taken altogether (‘omics green) with top 30 features from three individual models (see Methods). D-F) Discriminant features identified from gradient boosting classifiers significantly changed in the patient cohort compared to the healthy individuals. p-values were computed by Wilcoxon signed-rank test. p-value annotation legend: ns: $p > 0.05$, *: $0.01 < p <= 0.05$, **: $0.001 < p <= 0.01$, ***: $1e-04 < p <= 0.001$, ****: $p <= 1e-04$.

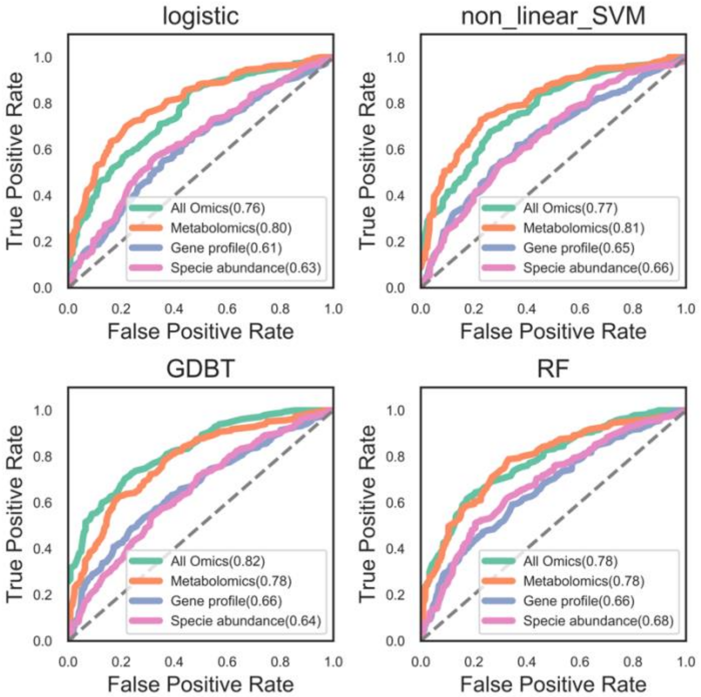


**Supplemental Figure 6. Multi-'omics models to classify the duration of ME/CFS (control vs. short-term vs. long-term).** As above, the performance of the classifiers using the area under the curve (AUC) was evaluated using 10 randomized and 10-fold cross-validations for each model: LASSO logistic regression, support vector machine (SVM), and random forest (RF) models. Models were designed based on species relative abundance (pink), KEGG gene profile (blue), or plasma metabolome (orange) individually, or taken all together ('omics green) with top 30 features from three individual models shown.

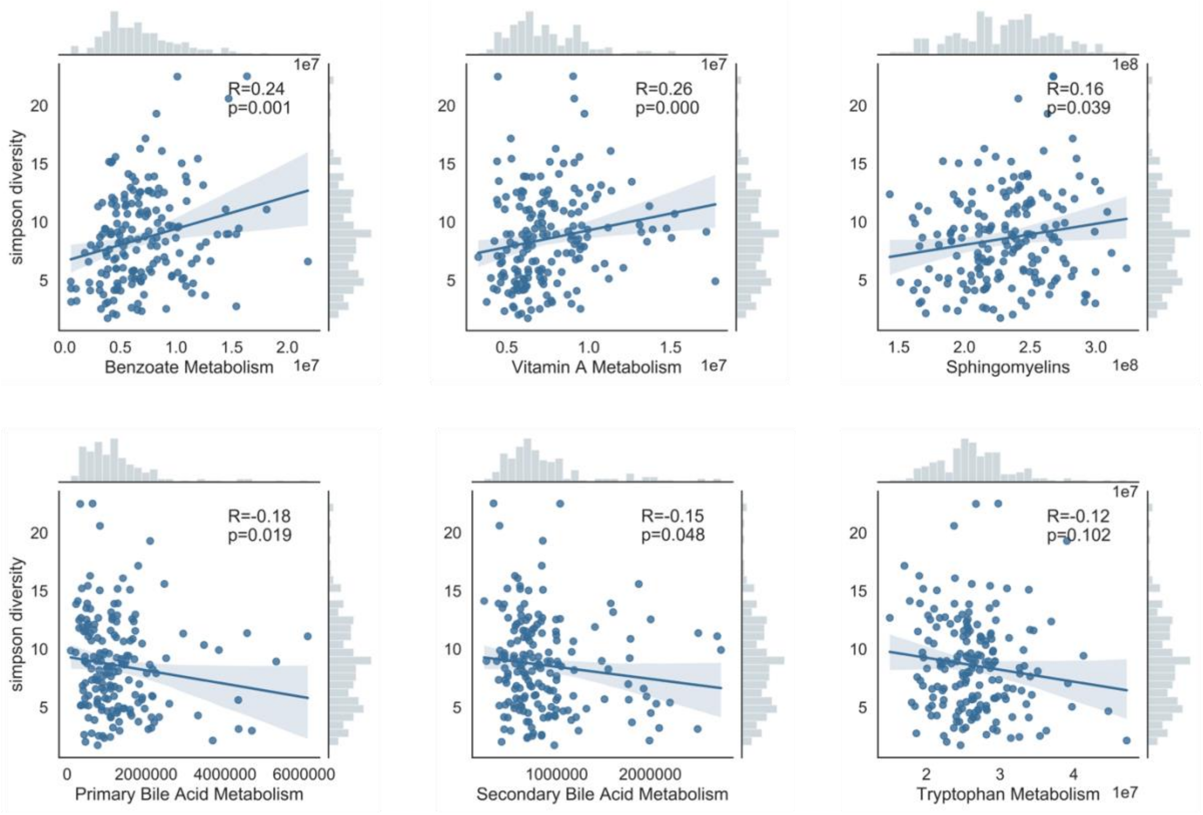


**Supplemental Figure 7. Plasma metabolic pathways are correlated with gut community structure in the cohort.** The gut microbiome community structure index was indicated by Simpson diversity and the relative abundance of the secondary metabolic pathway was indicated by the mean of all metabolites in the pathway. Each dot indicates one sample, and regression line and the confidence intervals shaded are shown. The data distributions are displayed in the top and right, respectively. The rho value and p-value were computed by the Spearman correlation.

### Supplemental Tables

**Supplementary Table 1** Demographic information of cohorts

**Supplementary Table 2** Clinical metadata summary

**Supplementary Table 3** Species relative abundances derived from MetaPhlAn3

**Supplementary Table 4** Predicted growth rate of species from GRiD

**Supplementary Table 5** KEGG gene reads count from metagenomic data

**Supplementary Table 6** Plasma metabolome raw data (peaks area-under-the-curve)

**Supplementary Table 7** Feature importance and annotation from multi-‘omics classification model

**Supplementary Table 8** Butyrate pathway: the statistics of key microbes and key enzymes in KEGG butanoate metabolism, and their correlations with key plasma metabolites.
